# Sampling motion trajectories during hippocampal theta sequences

**DOI:** 10.1101/2021.12.14.472575

**Authors:** Balázs B Ujfalussy, Gergő Orbán

**Affiliations:** Laboratory of Biological Computation, Inst. of Experimental Medicine, Budapest, Hungary; Laboratory of Neuronal Signalling, Inst. of Experimental Medicine, Budapest, Hungary; Computational Systems Neuroscience Lab, Wigner Research Center for Physics, Budapest, Hungary

## Abstract

Efficient planning in complex environments requires that uncertainty associated with current inferences and possible consequences of forthcoming actions is represented. **R**epresentation of uncertainty has been established in sensory systems during simple perceptual decision making tasks but it remains unclear if complex cognitive computations such as planning and navigation are also supported by probabilistic neural representations. Here we capitalized on gradually changing uncertainty along planned motion trajectories during hippocampal theta sequences to capture signatures of uncertainty representation in population responses. In contrast with prominent theories, we found no evidence of encoding parameters of probability distributions in the momentary population activity recorded in an open-field navigation task in rats. Instead, uncertainty was encoded sequentially by sampling motion trajectories randomly in subsequent theta cycles from the distribution of potential trajectories. Our analysis is the first to demonstrate that the hippocampus is well equipped to contribute to optimal planning by representing uncertainty.

## Introduction

Model-based planning and predictions are necessary for flexible behavior in a range of cognitive tasks. In particular, navigation is a domain that is ecologically highly relevant not only for humans but for rodents as well, which established a field for parallel investigation of the theory of planning, the underlying cognitive computations, and their neural underpinnings (***Hunt et al., 2021***). Importantly, predictions extending into the future have to cope with uncertainty coming from several different sources: uncertainty in the current state of the environment (our current location relative to a dangerous spot, the satiety of a predator or the actual geometry of the environment) and the availability of multiple future options when evaluating upcoming choices(***Glimcher, 2003; Redish, 2016***). Whether and how this planning-related uncertainty is represented in the brain is not known.

The hippocampus has been established as one of the brain areas critically involved in both spatial navigation and more abstract planning (***O’Keefe and Nadel, 1978; Miller et al., 2017***). **R**ecent progress in recording techniques and analysis methods largely contributed to understanding of the neuronal mechanisms underlying such computations (***Pfeiffer and Foster, 2013; Kay et al., 2020***).

A crucial insight gained about the neural code underlying navigation is that neuron populations in the hippocampus represent the trajectory of the animal on multiple time scales: Not only the current position of the animal can be read out at the behavioral time scale (***O’Keefe and Nadel, 1978; Wilson and McNaughton, 1993***), but also trajectories starting in the past and ending in the near future are repeatedly expressed on a shorter time scale at accelerated speed during individual cycles of the 6-10 Hz theta oscillation (theta sequences, ***Foster and Wilson 2007***, Fig. 1a). Moreover, features characteristic of planning can be identified in the properties of encoded trajectories, such as their dependence on the immediate context the animal is in, and on the span of the current run, future choices and rewards (***Johnson and Redish, 2007; Gupta et al., 2012; Wikenheiser and Redish, 2015; Tang et al., 2021; Zheng et al., 2020***). These data provide strong support for a computational framework where planning relies on sequential activity patterns in the hippocampus delineating future locations based on the current beliefs of the animal (***Stachenfeld et al., 2017; Miller et al., 2017***). Whether hippocampal computations also take into account the uncertainty associated with planning and thus the population activity represents the uncertainty of the encoded trajectories has not been studied yet.

**Figure 1:**
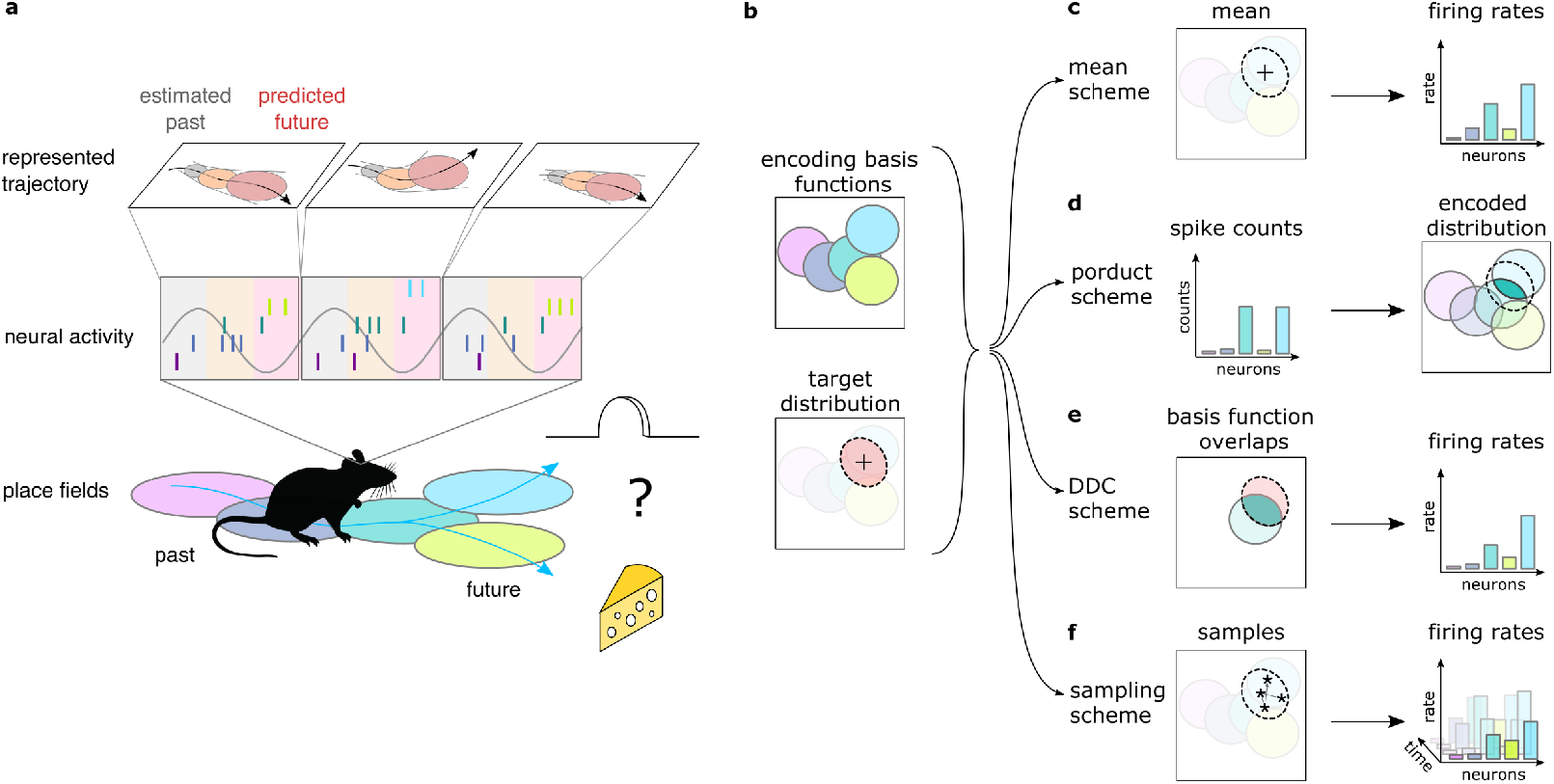
Theta sequences, uncertainty and variability. **a** Schematic showing the way hippocampal place cell activity represents possible trajectories in individual theta cycle during navigation. **b-f** Schemes for representing a target probability distribution (b, bottom) by the activity of a population of neurons each associated with an encoding basis function (related to their place field, Methods;b, top). **c** In the mean scheme the firing rate of the neurons (right, colored bars) is defined as the value of their basis functions (left, colored ellipses) at the mean of the target distribution (dotted ellipse). **d** In the product scheme the encoded distribution is proportional to the product of the basis functions of the active neurons. Note, that as opposed to the other schemes that map the target probability distribution to the firing rates, the product scheme defines the reverse mapping from the rates to the distribution. **e** In the DDC scheme the firing rate of each neuron is defined by the overlap between the target distribution and the basis function. **f** In the sampling scheme the firing rate of the neurons equals the value of their basis functions at the location sampled from the target distribution.

Neuronal representations of uncertainty have been extensively studied in sensory systems (***Ma et al., 2006; Orbán et al., 2016; Vértes and Sahani, 2018; Walker et al., 2020***). Schemes for representing uncertainty fall into three broad categories (Fig. 1d-f). In the first two categories (*product* and Distributed Distributional Code, *DDC*, schemes) the firing rate of a population encodes a complete probability distribution over spatial locations instantaneously at any given time by representing the parameters of the distribution (***Wainwright and Jordan 2008***, Methods). In the *product scheme* the firing rate of neurons encode a probability distribution through taking the product of the tuning functions (basis functions, Methods) of the coactive neurons (***Ma et al. 2006; Pouget et al. 2013***, Fig. 1d). In contrast, in the *DDC scheme* a population of neurons represents a probability distribution by signalling the overlap between the distribution and the basis function of individual neurons (***Zemel et al. 1998; Vértes and Sahani 2018***, Fig. 1e). In the third category, the *sampling scheme*, the population activity represents a single value sampled stochastically from the target distribution. In this case uncertainty is represented sequentially by the across-time variability of the neuronal activity (***Fiser et al. 2010***, Fig. 1f). These coding schemes provide a firm theoretical background to investigate the representation of uncertainty in hippocampus. Importantly, all these schemes have been developed for static features, where the represented features do not change in time (***Ma et al. 2006; Orbán et al. 2016***; but see ***Kutschireiter et al. 2017***). In contrast, trajectories represented in the hippocampus encode the temporally changing position of the animal. Here we extend the coding schemes to be able to accommodate the encoding of uncertainty associated with dynamic motion trajectories and investigate their neuronal signatures in rats while navigating an open environment.

In the present paper we propose that the hippocampus is performing probabilistic inference in a model that represents the temporal dependencies between spatial locations. Using a computational model, we demonstrate that key features of the hippocampal single neuron and population activity are compatible with representing uncertainty of motion trajectories in the population activity during theta sequences. Further, by developing novel computational measures, we pitch alternative schemes of uncertainty representation against each other and demonstrate that hippocampal activity does not show the hallmarks of schemes encoding probability distributions instantaneously. Instead, we demonstrate that the large and structured trial to trial variability between subsequent theta cycles is consistent with stochastic sampling from potential future trajectories but not with a scheme ignoring the uncertainty by representing only the most likely trajectory. These results demonstrate that the brain employs probabilistic computations not only in sensory areas during perceptual decision making but also in associative cortices during naturalistic, high-level cognitive processes.

## Results

### Neural variability increases within theta cycle

A key insight of probabilistic computations is that during planning uncertainty increases as trajectories proceed into more distant future (***Murphy, 2012; Sezener et al., 2019***). As a consequence, if planned trajectories are encoded in individual theta sequences, the uncertainty associated with the represented positions increases within a theta cycle (Fig. 1a). This systematic change in the uncertainty of the represented position during theta cycles is a crucial observation that enabled us to investigate the neuronal signatures of uncertainty during hippocampal theta sequences. For this, we analyzed a previously published dataset (***Pfeiffer and Foster, 2013***). Briefly, rats were trained to collect food reward in a 2×2 m large open arena from one of the 36 uniformly distributed food wells alternating between random forging and spatial memory task (***Pfeiffer and Foster, 2013***). Goal directed navigation in an open arena requires continuous monitoring and online correction of the deviations between the intended and actual motion trajectories. While sequences during both sharp waves and theta oscillations have been implicated in planning, here we focused on theta sequences as they are more strongly tied to the current position and thus averaging over many thousands of theta cycles can provide the necessary statistical power to identify the neuronal correlate of uncertainty representation.

Activity of hippocampal neurons was recorded by 20 tetrodes targeted to dorsal hippocampal area CA1 (***Pfeiffer and Foster, 2013***). Individual neurons typically had location-related activity (Fig. 2b, Fig. S6a), but their spike trains were highly variable (***Skaggs et al., 1996; Fenton and Muller, 1998***) (Fig. 2a). We used the empirical tuning curves (i.e., place fields) and a Bayesian decoder (***Zhang et al., 1998***) to estimate the represented spatial position from the spike trains of the recorded population in overlapping 20 ms time bins. Theta oscillations partition time into discrete segments and analysis was performed in these cycles separately (Fig. 2c). Despite the large number of recorded neurons (68–242 putative excitatory cells in 8 sessions from 4 rats), position decoding had a limited accuracy (Fisher lower bound on the decoding error in 20 ms bins: 16–30 cm vs. typical trajectory length ∼ 20 cm). Yet, in high spike count cycles we could approximately reconstruct the trajectories encoded in individual theta cycles (Fig. 2c). We then compared the reconstructed trajectories to the actual trajectory of the animal. We observed substantial deviation between the decoded trajectories and the motion trajectory of the animal: decoded trajectories typically started near the actual location of the animal and then proceeded forward (***Foster and Wilson, 2007***) often departing in both directions from the actual motion trajectory (***Kay et al. 2020***; Fig. 2c).

**Figure 2:**
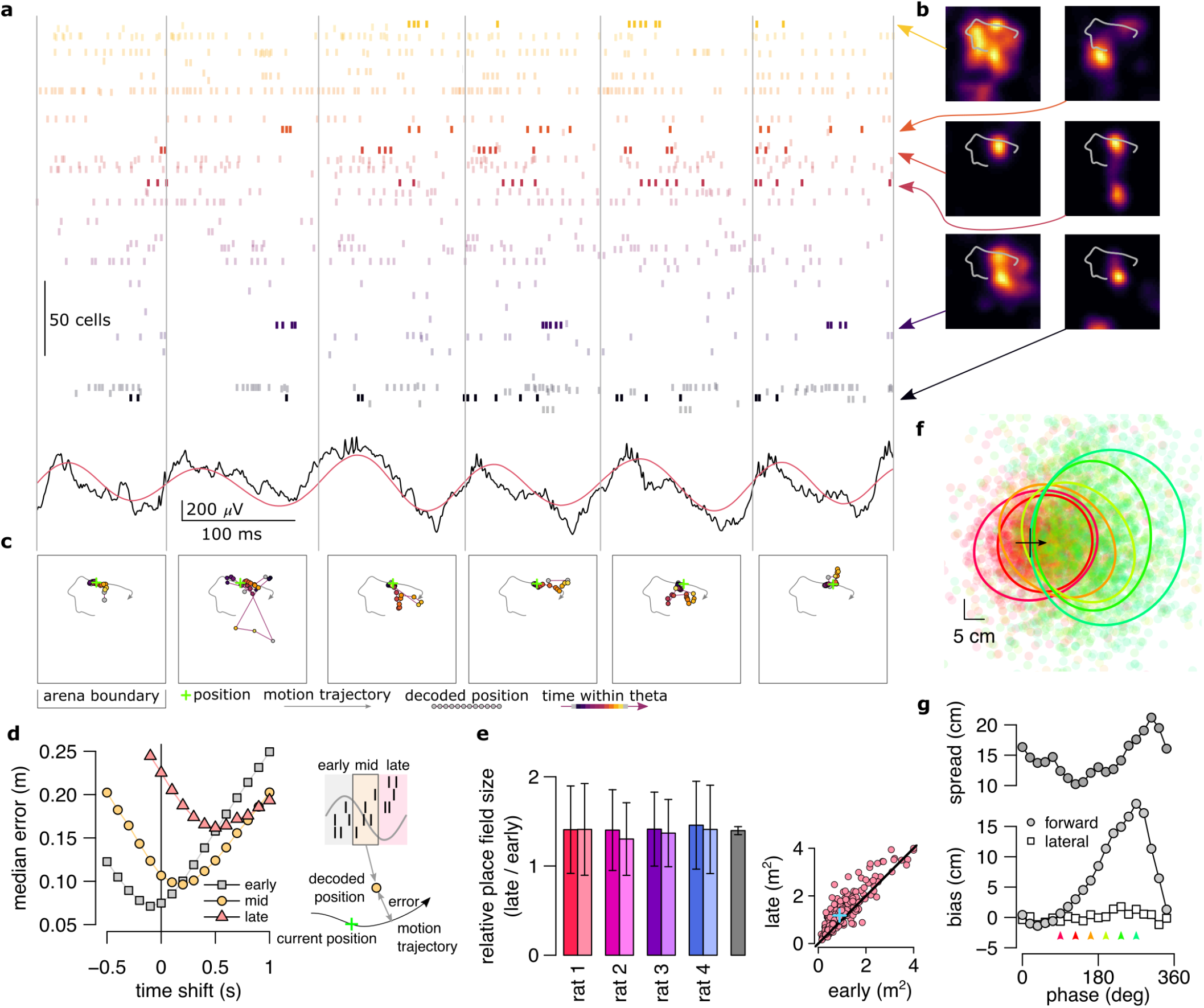
Theta sequences, uncertainty and variability. **a** Example spiking activity of 250 cells (top) and raw (black) and theta filtered (red) local field potential (bottom) for 6 consecutive theta cycles (vertical lines). **b** Place fields of 6 selected cells in a 2×2 m large open arena. Grey line indicates the motion trajectory during the run episode analysed in a-c. **c** Position decoded in overlapping 20 ms time bins (circles) during the 6 theta cycles shown in a. Time within theta cycle is color coded, grey indicates bins on the border of two cycles. Grey line shows the motion trajectory during the run episode, green crosses indicate the location of the animal in each cycle. **d** Decoding error for early, mid and late phase spikes (inset, top) calculated as a function of the time shift of the animal’s position along its motion trajectory (inset, bottom). For the analysis in panels d-e each theta cycle was divided into 3 parts with equal spike counts. **e** Relative place field size in late versus early theta spikes for the eight sessions (error bars: SD over cells). Grey bar: average and SD over all sessions. Inset: Place field size (ratemap area above 10% of the max firing rate) estimated from late vs. early theta spikes in an example session (individual dots correspond to individual place cells, blue cross: median). Only putative excitatory cells are included. To estimate the ratemaps, we shifted the reference positions with A*r* that minimised decoding error (see panel d). **f** Decoded positions (dots, in 20 ms bins) relative to the current position and motion direction (cross), and 0.5 confidence interval (CI) ellipses for 6 different theta phases (color, as in panel g). **g** Bias (bottom, mean of the decoded positions) and spread (top, see Methods) of decoded positions as a function of theta phase for an example session. Panels d,f,g show data from theta cycles with the highest 10% of spike counts.

To systematically analyse how this deviation depends on the theta phase we sorted spikes into three classes (early, mid and late). For any given class we decoded position from the spikes and compared it to the position of the animal shifted forward and backward along the motion trajectory. Time shift dependence of the accuracy of the decoders reveals the most likely portion of the trajectory the given class encodes (Fig. 2d, see also Fig. S4). For early spikes, the minimum of the average decoding error was shifted backward by ∼100 ms, while for late spikes +500 ms forward shift minimised the decoding error. These systematic biases indicate that, theta sequences tended to start slightly behind the animal and extended into the near future (***Foster and Wilson, 2007; Gupta et al., 2012***).

Further, the minima of the decoding errors showed a clear tendency: position decoded from later phases displayed larger average deviation from the motion trajectory of the animal (Fig. 2d). At the single neuron level, increased deviations at late theta phases resulted in the expansion of place fields (***Skaggs et al., 1996***): Place fields were more compact when estimated from early-phase spikes than those calculated from late-phase activity (Fig. 2e). At the population level, larger deviation between the motion trajectory and the late phase-decoded future positions can be attributed to the increased variability in encoded possible future locations. Indeed, when we aligned the decoded locations relative to the current location and motion direction of the animal, we observed that the spread of the decoded locations increased in parallel with the forward progress of their mean within theta cycles (Fig. 2f-g). Taken together, this analysis demonstrated that the cycle-to-cycle variability of the encoded trajectory is larger at the end of the theta cycle when encoding more uncertain future locations than at the beginning of the cycle when representing the past position.

The observed increase of the cycle-to-cycle variability could be a direct consequence of encoding variable 2-dimensional trajectories (Fig. S4a-b) or it may be a signature of the neuronal representation of uncertainty. In the next sections we set up a synthetic dataset to investigate the theta sequences and cycle-to-cycle variability in the four alternative coding schemes.

### Synthetic data: testbed for discriminating the encoding schemes

To analyse the distinctive properties of the different coding schemes and to develop specific measures capable of discriminating them, we generated a synthetic dataset in which both the behavioral component and the neural component could be precisely controlled. The behavioural component was matched to the natural movement statistics of rats during navigating a 2D environment. The neural component was constructed such that it could accommodate the alternative representational schemes for prospective representations during navigation.

Our synthetic dataset had three consecutive levels. First, we simulated a planned trajectory for the rat in a two dimensional open arena by allowing smooth changes in the speed and motion direction (Methods, Fig. S1a-b). Second, similar to the real situation, the simulated animal did not have access to its true position, *x*_*n*_, in theta cycle *n*, but had to infer it from the sensory inputs it had observed in the past, *y*_1:*n*_ (Methods). To perform this inference and predict its future position, the animal used a model of its own movements in the environment. In this dynamical generative model the result of the inference is a posterior distribution over possible trajectories starting *n*_*p*_ steps back in the past and ending *n*_*f*_ steps forward in the future. To generate the motion trajectory of the animal noisy motor commands were generated based on the difference between its planned trajectory and its inferred current position (Methods, Fig. S1a-b, Fig. S2). The kinematics was matched between simulation and experimental animals (Fig. S2).

Third, in our simulations the hippocampal population activity encoded an inferred trajectory at an accelerated speed in each theta cycle such that the trajectory started in the past at the beginning of the simulated theta cycle and led to the predicted future states (locations) by the end of the theta cycle (Fig. 3a-c). To match the size of the synthetic and experimental data, we simulated the activity of 200 hippocampal pyramidal cells (*place cells*). Firing rates of pyramidal cells depended on the encoded spatial location using either of the 4 different coding schemes (Methods): In the *mean* code, the population encoded the single most likely trajectory without representing uncertainty. In *product* and *DDC* schemes, a snapshot of the population activity at any given theta phase encoded the estimated past or predicted future part of the trajectory in the form of a probability distribution (Methods). Finally, in the sampling scheme, in each theta cycle a single trajectory was sampled stochastically form the distribution of possible trajectories (Fig. S1). The posterior distribution was updated in every ∼100 ms matching the length of theta cycles. Importantly, all of the 4 encoding schemes yielded single neuron and population activity dynamics consistent with the majority of known feature of hippocampal population activity including spatial tuning, phase precession (Fig. 3d) and theta sequences (Fig. 3e, see also Fig. S3).

**Figure 3:**
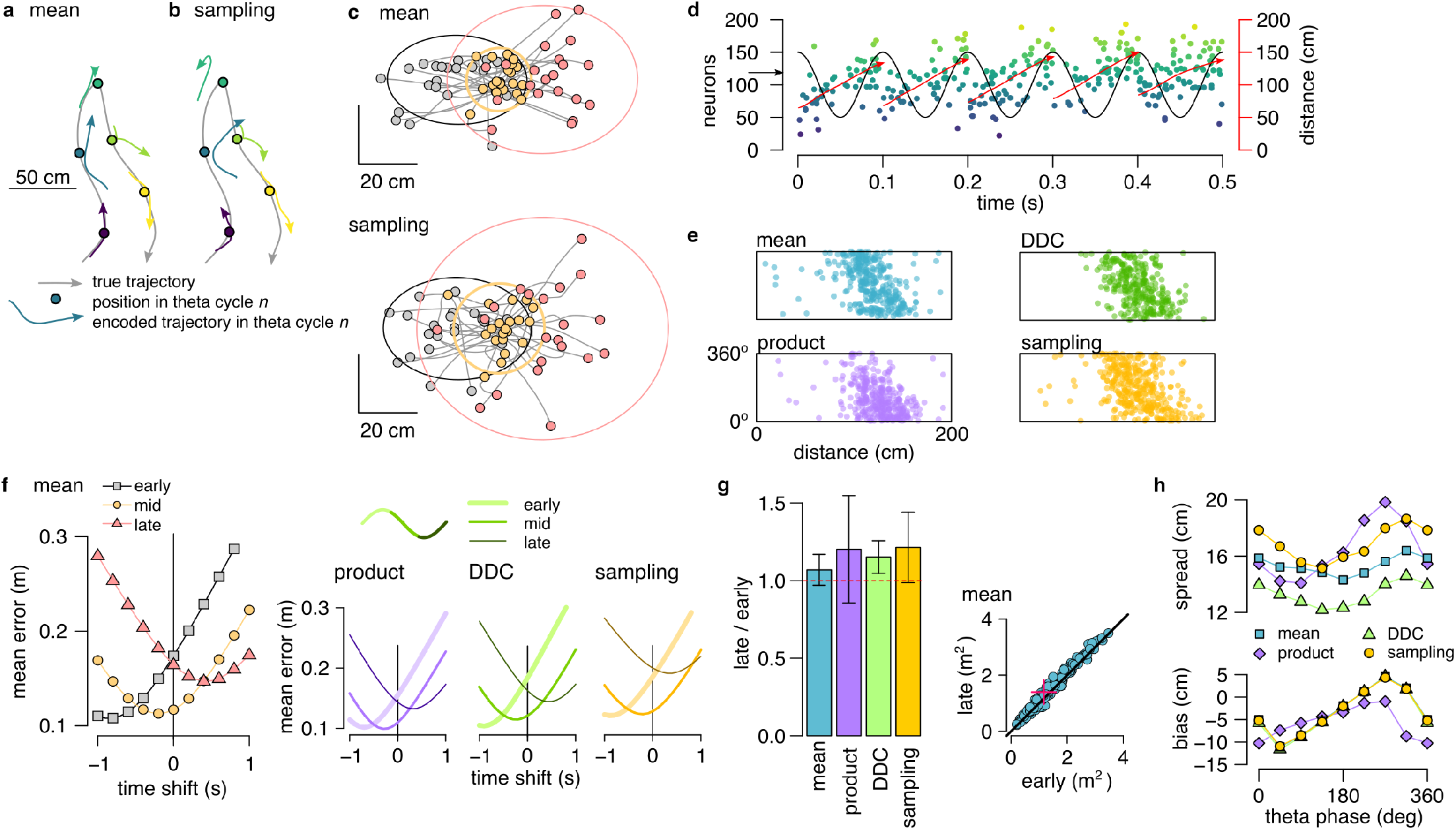
Theta sequences in simulated data. **a** Motion trajectory of the simulated rat (grey, 10 s) together with its own inferred and predicted most likely (mean) trajectory segments (colored arrows) in five locations separated by 2 s (filled circles). These trajectories were represented in theta cycles using one of the four alternative schemes. **b** Same as panel A, but trajectories sampled from the posterior distribution. **c** Example represented trajectories aligned to the actual position and direction of the simulated animal in the mean (top) and in the sampling (bottom) scheme. Ellipses indicate 50% CI of all theta cycles. Color code indicates start (grey), mid (yellow) and end (pink) of the trajectories. **d** Simulated activity of 200 place cells sorted according to the location of their place fields on a linear track (200 × 10 cm) during an idealised 10 Hz theta oscillation using the mean encoding scheme. Red lines show the 1-dimensional trajectories represented in each theta cycle. Note that the overlap between trajectories is larger here than in panels a-b, because, for clarity, only trajectories at every 20th theta cycle is shown there. **e** Theta phase of spikes of an example simulated neuron (arrow in panel d) as a function of the animal position in the four coding schemes. **f** Decoding error from early, mid and late phase spikes (highest 5% spike count cycles) as a function of the time shift of the simulated animal’s position in a mean, product, DDC and sampling schemes. **g** Relative place field size in late versus early theta spikes for the four different models (error bars: SD over 200 cells). Inset: place field size estimated from late vs. early theta spikes in the mean scheme. Median is indicated with red cross. **h** Decoding bias (bottom) and spread (top) as a function of theta phase for the four different encoding models (n_sp_ > 0% -product model is biased otherwise).

After generating the synthetic datasets we investigated how positional information is represented during theta sequences in each of the four alternative coding schemes. We decoded the synthetic population activity in early, mid and late theta phases and compared the estimated position with the actual trajectory of the simulated animal. The deviation between the decoded position and the motion trajectory increased throughout the theta cycle irrespective of the coding scheme (Fig. 3f). Moreover, place fields were significantly larger when estimated from late than early-phase spikes in all four representational schemes (Fig. 3g, P *<* 10^−16^ for each scheme). Finally, when aligned to the current position and motion direction of the simulated animal, the spread of the decoded locations increased with the advancement of their mean within theta cycles in all four coding schemes (Fig. 3h).

Our analyses thus confirmed that the increased variability of the hippocampal neural activity at late versus early theta phase is consistent with encoding trajectories in a dynamical model of the environment irrespective of the representation of the uncertainty. These synthetic datasets provide a functional interpretation of the hippocampal population activity during theta oscillations and offer a unique testbed for contrasting different probabilistic encoding schemes. In the following sections we will identify hallmarks for each of the representational schemes of uncertainty and use these hallmarks to discriminate them through the analysis of neural data (Fig. 4a).

**Figure 4:**
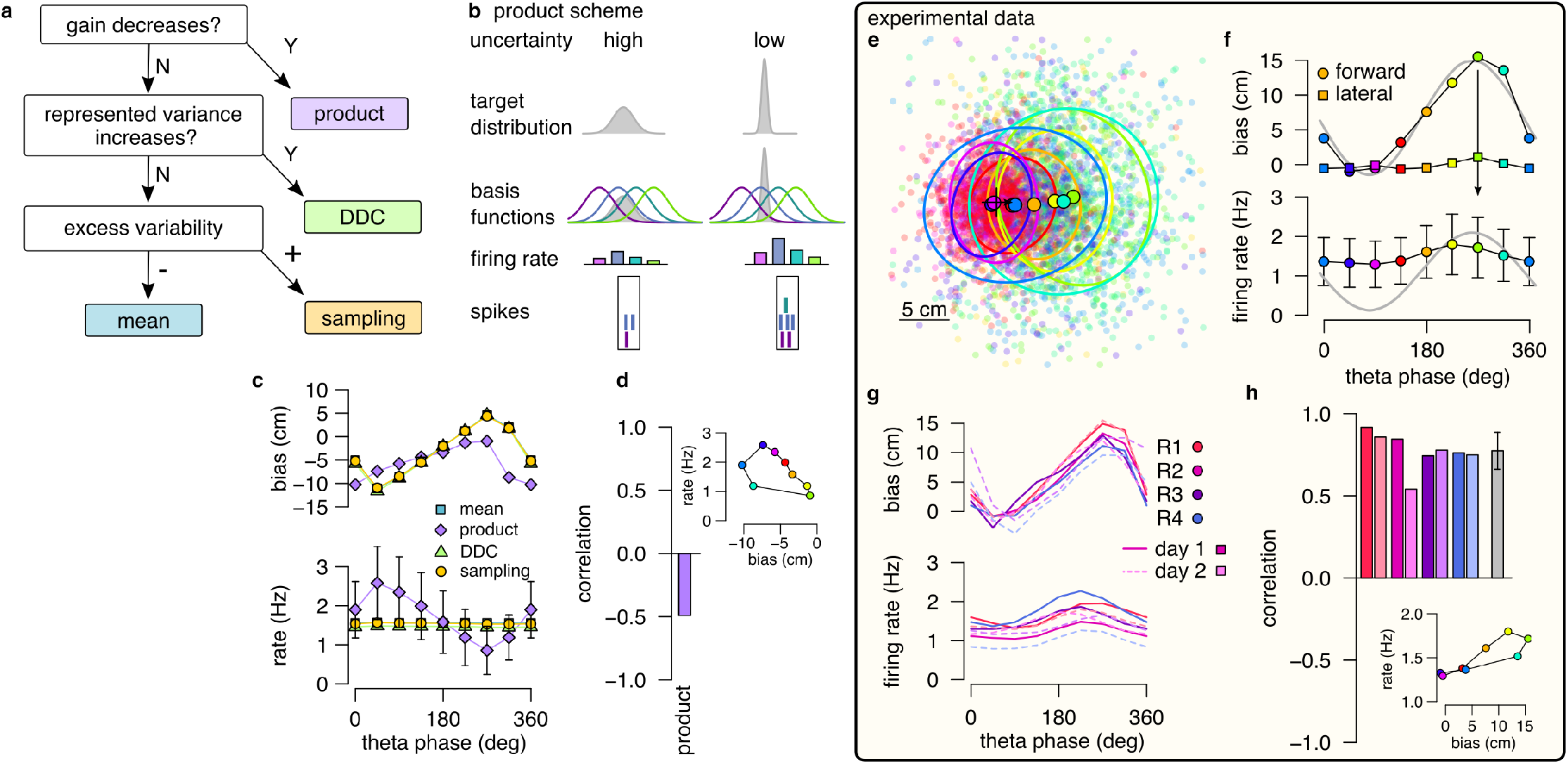
Product scheme: population gain decreases with uncertainty. **a** Decision-tree for identifying the representational scheme. **b** Schematic of encoding a high-uncertainty (left) and a low-uncertainty (right) target distribution using the product scheme with 4 neurons. The SD is represented by the gain of the population. **c** Population firing rate (bottom) and forward decoding bias (top) as a function of theta phase for the four schemes in synthetic data. Only the product scheme predicts a systematic change in the firing rate. **d** Correlation between firing rate and forward bias for the product scheme. Inset: Firing rate as a function of forward bias in the product scheme. Color code is the same as in f. **e** Decoded position relative to the location and motion direction of the animal (black arrow) at eight different theta phases in an example recording session. Filled circles indicate mean, ellipses indicate 50% CI of the data. (Similar to Fig. 3g, but with 120^°^and not 20 ms binning.) **f** Forward decoding bias of the decoded position (top) and population firing rate (bottom) as a function of theta phase in an example recording session. Grey line in top and bottom show cosine fit to the forward decoding bias. **g** Decoding bias (top) and firing rate (bottom) for all animals and sessions (line type). **h** Correlation between firing rate and forward bias for all recorded sessions. Grey bar: mean and SD across the eight sessions. Inset: Firing rate as a function of forward decoding bias in an example session. Bias in e-g was calculatd using the 5% highest spike count theta cycles. Population firing rate plots show average over all cycles.

### Testing the product scheme: gain

To discriminate the product scheme from other representations we capitalise on the specific relationship between response intensity of neurons and uncertainty of the represented variable. In a product representation, a probability distribution over a feature, such as the position, is encoded by the product of the neuronal basis functions (Methods). When the basis functions are localised, as in the case of hippocampal place fields, the width of the encoded distribution tends to decrease with the total number of spikes in the population (Fig. 4b, Supplemental note, Fig. S6). There-fore we propose using the systematic decrease of the population firing rate (*gain*) with increasing uncertainty as a hallmark of the product scheme.

We first tested for the specificity of the co-modulation of the population gain with uncertainty to the product scheme and compared gain modulation in the four different coding schemes in our synthetic dataset. In each of the coding schemes, we identified the theta phase with the maximal uncertainty by the maximal forward bias in encoded positions. For this, we decoded positions from spikes using eight overlapping 120^°^windows in theta cycles and defined the end of the theta cycles based on the maximum of the average forward bias (Fig. Fig. 4C top). Then we calculated the average number of spikes in a given 120^°^window as a function of the theta phase. We found that the product scheme predicted a systematic, ∼3-fold modulation of the firing rate within the theta cycle (Fig. 4c, bottom). The peak of the firing rate coincided with the theta phase encoding backward positions, when the uncertainty is minimal and the correlation between the firing rate and the forward bias was negative (Fig. 4d). The three other coding schemes did not constrain the firing rate of the population to represent probabilistic quantities, and thus the firing rate was independent of the theta phase or the encoded uncertainty.

After demonstrating the specificity of uncertainty-related gain modulation to the product scheme, we returned to the experimental dataset and applied the same analysis to neuronal activity recorded from freely navigating rats. We first decoded the spikes during theta oscillation falling in eight overlapping 120^°^window and aligned the decoded locations relative to the animals’ position and motion direction. We confirmed that the encoded location varied systematically within the theta cycle from the beginning towards the end of the theta cycle both when considering all theta cycles (Fig. S4) or when constraining the analysis to theta cycles with the highest 5% spike count (Fig. 4e-f; cf. Fig. 2f-g). Maximal population firing rate coincided with the end of the theta sequences which correspond to future positions characterized by the highest uncertainty (Fig. 4f). Thus, a positive correlation emerged between the represented uncertainty and the population gain (Fig. 4h). This result was consistent across recording sessions and animals (Fig. 4g-h) and was also confirmed in an independent dataset where rats were running on a linear track (Fig. S5a-d; ***Grosmark et al. 2016***).

This observation is in sharp contract with the prediction of the product encoding scheme where the maximum of the firing rate should be near the beginning of the theta sequences (Fig. 4c). The other encoding schemes are neutral about the theta modulation of the firing rate, and therefore they are all consistent with the observed small phase-modulation of the firing rate.

### Testing the DDC scheme: diversity

Next, we set out to discriminate the DDC scheme from the mean and the sampling schemes. In the DDC scheme, neuronal firing rate represents the overlap of the basis functions with the encoded distribution (Fig. 1e; ***Zemel et al. 1998; Vértes and Sahani 2018***). Naturally, in this scheme the diversity of the co-active neurons increases with the variance of the encoded distribution (Fig. 5a). When the encoded variance is small, only neurons with overlapping basis functions will fire synchronously. Conversely, a large diversity of neurons become co-active when encoding a distribution of high variance (Fig. 5a). Therefore the width (standard deviation, SD) of the encoded distribution can be decoded from the population activity and we propose to use the increase of decoded SD with increasing uncertainty as a hallmark of DDC encoding.

**Figure 5:**
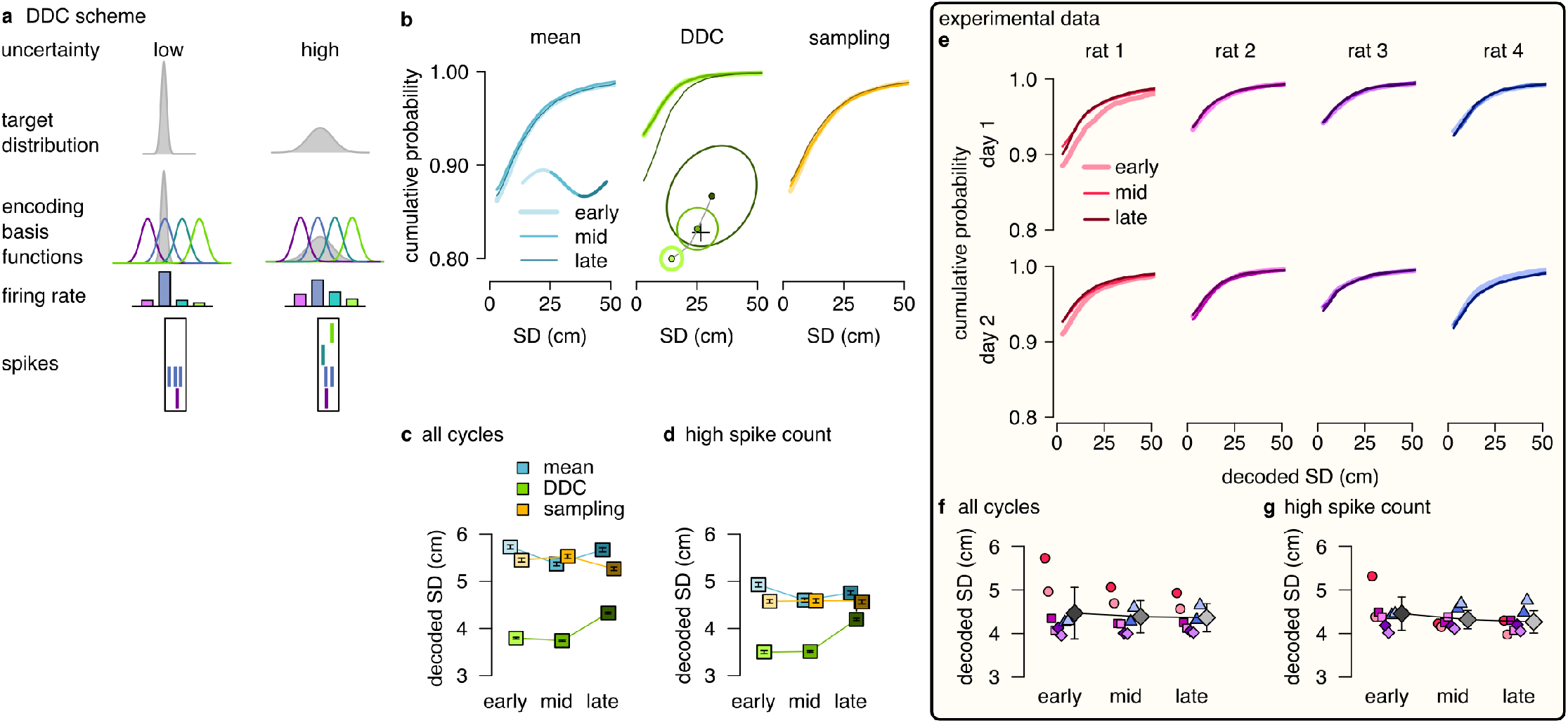
DDC scheme: diversity increases with uncertainty. **a** Schematic of encoding a narrow (left) and a wide (right) distribution with spike-based DDC using 4 neurons. Intuitively, the SD is represented by the diversity of the co-active neurons. **b** Cumulative probability of the decoded SD of the represented distribution at different theta phases for the mean, DDC and sampling schemes in the simulated dataset. **c** Mean and SE of decoded SD as a function of theta phase for the different schemes in the simulated dataset. Only the DDC code predicts a slight, but significant increase in the decoded SD at late theta phases. **d** Same as c calculated from theta cycles with higher than median spike count. **e** Cumulative probability of the decoded SD from spikes in early, mid and late phase of the theta cycles (all cycles) for the analysed sessions. **f-g** Mean of the decoded SD for each animal from early, mid and late theta spikes using all theta cycles (f) or theta cycles with higher than median spike count (g). Symbols show mean and SD across animals. See also Fig. S8 for similar analysis using the estimated encoding basis functions instead of the empirical tuning curves for decoding.

First, we turned to our synthetic dataset to demonstrate that diversity of population responses can be used to estimate SD in a DDC code and that systematic changes in decoded SD are specific to this scheme. In each of the three remaining coding schemes (mean, DDC and sampling) we divided the population activity to three distinct windows relative to the theta cycle (early, mid and late). We decoded the mean and the SD of the encoded distribution of trajectories (Fig. 5b inset) using the empirical tuning curves (Meethods). We found a systematic and significant increase in the decoded SD in the DDC scheme from early to late theta phases (one-sided, two sample Kolmogorov-Smirnov test, *P* = 1.3 × 10^−16^), whereas the decoded SD was independent of the theta phase for the mean and the sampling schemes (KS test, mean scheme: *P* = 0.98, sampling scheme: *P* = 0.95, Fig. 5b-c). This result was robust against using the theta cycles with higher than median spike count for the analysis (Fig. 5d, mean: *P* = 0.99, DDC: *P* = 8.2 × 10^−12^, sampling: *P* = 0.93) or against using the estimated basis functions instead of the empirical tuning curves for decoding (Fig. S8a-b; Supplemental note). Thus, our analysis of synthetic data demonstrated that the decoded SD is a reliable measure to discriminate the DDC scheme from sampling or mean encoding.

After testing on synthetic data, we repeated the same analysis on the experimental dataset. We divided each theta cycle into three windows of equal spike counts (early, mid and late) and decoded the mean and the SD of the encoded trajectories from the population activity in each window. We found that the decoded SDs had a nearly identical distribution at the three theta phases for all recording sessions (Fig. 5e). The mean of the decoded SD did not change significantly or consistently across the recording session neither when we analysed all theta cycles (early vs. late, KS test *P >* 0.62 for all sessions, Fig. 5e-f) nor when we constrained the analysis to the half of the cycles with higher than median spike count (KS test, *P >* 0.7 for all sessions, Fig. 5g) or when we used the estimated encoding basis functions instead of the empirical tuning curves for decoding (Fig. S8c-d). We obtained similar results in a different dataset with rats running on a linear track (Fig. S5e-i; ***Grosmark et al. 2016***). We conclude that there are no signatures of DDC encoding scheme in the hippocampal population activity during theta oscillation. Taken together with the findings on the product scheme, our results indicate that hippocampal neurons encode individual trajectories rather than entire distributions during theta sequences.

### Testing the sampling scheme: excess variability

Both the MAP scheme and the sampling scheme represent individual trajectories but only the sampling scheme is capable of representing uncertainty. Therefore a critical question concerns if the two schemes can be distinguished based on the population activity during theta sequences. Sampling-based codes are characterised by large and structured trial-to-trial neuronal variability (***Orbán et al., 2016***). Our results showed a systematic increase in the variability of the decoded location at phases of theta oscillation that correspond to later portions of the trajectory associated with higher uncertainty. This parallel increase of variability and uncertainty could be taken as evidence towards the sampling scheme. However, we demonstrated that the systematic theta phase-dependence of the neural variability and the variability of the encoded trajectories is a general feature of predictions in a dynamical model characterising both the sampling and the MAP schemes (Fig. 2-3). In the sampling scheme, the cycle-to-cycle variability is further increased by the sampling variance, i.e. the stochasticity of the represented trajectory, such that the magnitude of excess variance is proportional to uncertainty. In order to discriminate sampling from the mean scheme we designed a measure, excess variability, that can identify the additional variance resulting from sampling. For this, we partitioned the variability of encoded trajectories such that the full magnitude of variability across theta cycles (cycle-to-cycle variability, *CCV*) is compared to the amount of variability expected from the animal’s own uncertainty (trajectory encoding error, *TEE*, Fig. 6a, Methods).

**Figure 6:**
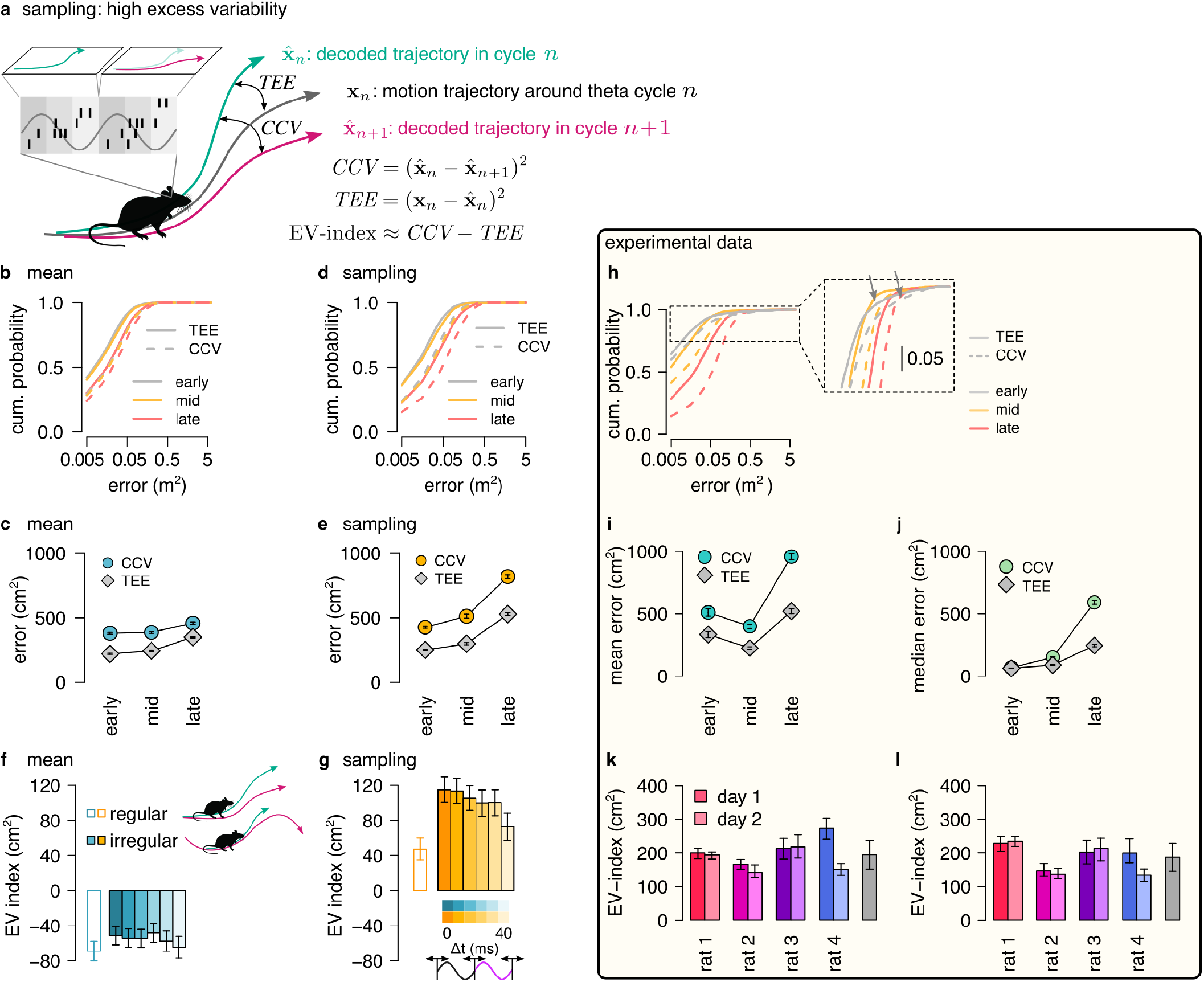
Sampling scheme: excess variability increases with uncertainty. **a** To discriminate sampling from mean encoding we defined the EV-index which measures the magnitude of cycle-to-cycle variability (*CCV*) relative to the trajectory encoding error *(TEE)*. **b** Cumulative distribution of *CCV* (dashed) and *TEE* (solid) for early, mid and late theta phase (colors) in the mean scheme using simulated data. Note the logarithmic x axis. **c** Mean *CCV* and *TEE* calculated from early, mid and late phase spikes in the mean scheme. **d-e** Same as b-c for the sampling scheme. **f-g** The EV-index for the mean (f) and sampling (g) schemes with simulating regular or irregular theta-trajectories (left inset) and applying various amount of jitter for segmenting the theta cycles (color code, right inset). Error bars show SE. **h** Cumulative distribution of *CCV* (dashed) and *TEE* (solid) for early, mid and late theta phase (colors) for an example recording session. Note the logarithmic x axis. Arrows in inset highlight atypically large errors occurring mostly at early theta phase. **i-j** Mean (i) and median (j) of *CCV* and *TEE* calculated from early, mid and late phase spikes for the session shown in h. **k-l** EV-index calculated for all analysed sessions (color) and across session mean and SD (grey) using the mean (k) or the median (l) across theta cycles. Error bars show SE in k and 5% and 95% confidence interval in l. P-values are shown in Table 2. Here we analysed only theta cycles with higher than median spike count. See Fig. S9 for similar analysis including all theta cycles.

We assessed excess variability in our synthetic dataset using either the sampling or the mean representational schemes, i.e. encoding either sampled trajectories or mean trajectories. Specifically, we decoded the population activity in three separate windows of the theta cycle (early, mid and late) using a standard static Bayesian decoder and computed the difference between the decoded locations across subsequent theta cycles (cycle-to-cycle variability, *CCV*) and the difference between the decoded position and the true location of the animal (trajectory encoding error, *TEE*). As expected, both *TEE* and *CCV* increased from early to late theta phase both for mean (Fig. 6b-c) and sampling codes (Fig. 6d-e, Methods). Our analysis confirmed that it is the magnitude of the increase that is the most informative of the identity of the code: the increase of *CCV* is more intense within theta cycle than *TEE* in the case of sampling (Fig. 6e) whereas increase of the *TEE* is larger in the mean encoding (Fig. 6c, Methods). We quantified this difference using the *excess variability index* (EV-index). The EV-index was negative for mean (Methods, Fig. 6f) and positive for sampling schemes (Fig. 6g). To test the robustness of the EV-index against various factors influencing the cycle-to-cycle variability we analyzed potential confounding factors. First, to compensate for the potentially large decoding errors during low spike count theta cycles, we calculated the EV-index both using all theta cycles (Fig. S9e-f) or only theta cycles with high spike count (Fig. 6f-g). Second, we varied the speed and the length of the encoded trajectories (irregular vs. regular trajectories, Fig. 6f inset, Methods). Third, we introduced a jitter to the boundaries of the theta cycles in order to model our uncertainty about cycle boundaries in experimental data (jitter 0-40 ms, Fig. 6g inset, Methods). We found that the EV-index was robust against these manipulations, reliably discriminating sampling based codes from mean codes across a wide range of parameters. Thus, EV-index can distinguish sampling related excess variability from variability resulting from other sources.

Finally we repeated the same analysis on the dataset recorded from rats exploring the 2D spatial arena. We calculated cycle-to-cycle variability and trajectory encoding error and found that although typically both the error and the variability increased during theta, their distributions had high positive skewness (Fig. 6h). The outliers preferentially occurred at early theta phase (grey arrows in Fig. 6h), when they increased the mean of the distribution above its mid-theta value (Fig. 6i). The median was more robust against the effect of the outliers (Fig. 6j). Therefore we calculated the EV-index using both the mean and the median across all theta cycles (Fig. S9j-k) or including only high spike count cycles (Fig. 6k-l). We found that the EV-index was consistently and significantly positive for all recording sessions. This analysis supports that the large cycle-to-cycle variability of the encoded trajectories during hippocampal theta sequences is consistent with random sampling from the posterior distribution of possible trajectories in a dynamic generative model.

## Discussion

In this paper we demonstrated that the statistical structure of the sequential neural activity during theta oscillation in the hippocampus is consistent with repeatedly performing inference and predictions in a dynamical generative model. Further, we have established a framework to directly contrast competing hypotheses about the way probabilistic inference can be performed in neural populations. Importantly, new measures were developed to dissociate alternative coding schemes and to identify their hallmarks in the population activity. **R**eliability and robustness of these measures was validated on synthetic data with characteristics closely matching the experimental conditions. Our analysis demonstrated that the neural code in the hippocampus shows hallmarks of efficient planning, that is it is capable of representing information about uncertainty. Specifically, our analysis has shown that the hippocampal population code reflects the signature of sampling possible motion trajectories but could not identify the hallmarks of alternative proposed schemes for coding uncertainty.

### Planning and dynamical models

Hippocampal activity sequences both during theta oscillation and sharp waves have been implicated in planning. During sharp waves, sequential activation of place cells can outline extended trajectories up to 10 m long (***Davidson et al., 2009***), a mechanism suitable for selecting the correct path leading to goal locations (***Pfeiffer and Foster, 2013; Mattar and Daw, 2018***). During theta oscillation, sequences typically cover a few tens of centimeters (Fig. 2; ***Wikenheiser and Redish 2015***) and are thus more suitable for surveilling the immediate consequences of the imminent actions. Monitoring future consequences of actions at multiple time scales is a general strategy for animals, humans and autonomous artificial agents alike (***Neftci and Averbeck, 2019***).

In our model the dominant source of uncertainty represented by the hippocampal population activity was the stochasticity of the animal’s forthcoming choices. Indeed, it has been observed that before the decision point in a W-maze the hippocampus also represents hypothetical trajectories not actually taken by the animal (***Redish, 2016; Kay et al., 2020; Tang et al., 2021***). Here we generalised this observation to an open field task and found that monitoring alternative paths is not restricted to decision points but the hippocampus constantly monitors consequences of alternative choices. Disentangling the potential contribution of other sources of uncertainty, including ambiguous sensory inputs (***Jezek et al., 2011***) or unpredictable environments to hippocampal representations requires analysing population activity during more structured experimental paradigms (***Kelemen and Fenton, 2010; Miller et al., 2017***).

Although representation of alternative choices in theta sequences is well supported by data, it is not clear how much the encoded trajectories are influenced by the current policy of the animal. On one hand, prospective coding, reversed sequences during backward travel or the modulation of the sequences by goal location indicates that current context influences the content and dynamics of the sequences (***Frank et al., 2000; Johnson and Redish, 2007; Cei et al., 2014; Wikenheiser and Redish, 2015***). On the other hand, theta sequences represent alternatives with equal probability in binary decision tasks and the content of the sequences is not predictive about the future choice of the animal (***Kay et al., 2020; Tang et al., 2021***). Sampling from a relatively wide proposal distribution is a general motif utilised by several sampling algorithms (***MacKay, 2003***). Brain areas beyond the hippocampus, such as the prefrontal cortex, might perform additional computations on the array of trajectories sampled during theta oscillations, including rejection of the samples not consistent with the current policy (***Tang et al., 2021***). The outcome of the sampling process can have profound effect on future behavior as selective abolishment of theta sequences by pharmacological manipulations impairs behavioral performance (***Robbe et al., 2006***).

We found that excess variability in the experimental data was higher than that in the simulated data. In our model we sampled independent trajectories in each theta cycle but more efficient algorithms can further increase the difference between subsequent samples by preferentially selecting samples from the opposite lobes of the target distribution (***MacKay, 2003***) leading to generative cycling (***Kay et al., 2020***). Such effective sampling algorithms can increase excess variability. **R**ecurrent neural networks optimised for efficient sampling were shown to display strong oscillatory activity in the gamma band (***Echeveste et al., 2020***). Concurrent gamma and theta band activities are characteristics of hippocampus (***Colgin, 2016***), which indicates that network mechanisms underlying efficient sampling might be exploited by the hippocampus too. Efficient sampling of trajectories instead of static variables could necessitate multiple interacting oscillations where individual trajectories evolve during gamma cycles and alternative trajectories are sampled in theta cycles.

### Circuit mechanisms

**R**ecent theoretical studies established that recurrent neural networks can implement complex nonlinear dynamics (***Mastrogiuseppe and Ostojic, 2018; Vyas et al., 2020***) including sampling from the posterior distribution of a static generative model (***Echeveste et al., 2020***). External inputs to the network can efficiently influence the trajectories emerging in the network either by changing the internal state and initiating a new sequence (***Kao et al., 2021***) or by modulating the internal dynamics influencing the transition structure between the cell assemblies or the represented spatial locations (***Mante et al., 2013; Stroud et al., 2018***). We speculate that the recurrent network of the CA3 subfield could serve as a neural implementation of the dynamical generative model with inputs from the entorhinal cortex providing strong contextual signals selecting the right map and conveying landmark information necessary for periodic resets at the beginning of the theta cycles.

Although little is known about the role of entorhinal inputs to the CA3 subfield during theta sequences, inputs to the CA1 subfield show functional segregation consistent with this idea. Specifically, inputs to the CA1 network from the entorhinal cortex or from the CA3 region are activated dominantly at distinct phases of the theta cycle and are associated with different bands of gamma oscillations reflecting the engagement of different local micro-circuits (***Colgin et al., 2009; Schomburg et al., 2014***). The entorhinal inputs are most active at early theta phases when they elicit fast gamma oscillations (***Schomburg et al., 2014***) and these inputs might contribute to the reset of the sequences (***Fernández-Ruiz et al., 2017***). Initial part of the theta cycle showing transient backward sequences (***Wang et al., 2020***) could reflect the effect of external inputs resetting the local network dynamics. Conversely, CA3 inputs are coupled to slow gamma rhythms preferentially occurring at later theta phases, associated with prospective coding, and relatively long, temporally compressed paths (***Schomburg et al., 2014; Bieri et al., 2014***) potentially following the trajectory outlined by the CA3 place cells.

### Representations of uncertainty

Uncertainty representation implies that not only a best estimate of an inferred quantity is maintained in the population but properties of a full probability distribution can also be recovered from the activity. Machine learning provides two major classes of computational methods to represent these distributions: 1, instantaneous representations which rely on a set of parameters to encode a probability distribution; or 2, sequential representations that collect samples from the distributions (***MacKay, 2003***). Accordingly, theories of neural probabilistic computations fall into these categories: the product scheme and DDC instantaneously, while sampling sequentially represents uncertainty (***Pouget et al., 2013; Savin and Deneve, 2014***). Our analysis did not find evidence for representing a probability distributions instantaneously during hippocampal theta sequences. Instead, our data is consistent with representing a single location at any given time where uncertainty is encoded sequentially by the variability of the represented locations across time.

Importantly, it is not possible to accurately recover the represented position of the animal from the observed spikes in any of the coding schemes: one can only estimate it with a finite precision, summarised by a posterior distribution over the possible positions. Thus, decoding noisy neural activity naturally leads to a posterior distribution. However, this does not imply that it was actually a probability distribution encoded in the population activity (***Zemel et al., 1998; Lange et al., 2020***). Specifically, when a scalar variable *x* is encoded in the spiking activity *s*(*x*) of neurons using an exponential family likelihood function, then the resulting code is a linear probabilistic population code (***Ma et al., 2006***) (PPC). In fact, we used an exponential family likelihood function (Poisson) in our mean and sampling scheme, so these schemes belong, by definition, to the PPC family. However, the PPCs should not be confused with our product scheme where a reference distribution is encoded in the population activity instead of a single variable (***Lange et al., 2020***).

To test the product scheme, we used the population gain as a hallmark. We found that the gain varied systematically, but the variance was not consistent with the basic statistical principle, that on average uncertainty accumulates when predicting future states. An alternative test would be to estimate both the encoding basis functions and the represented distributions as proposed by ***Ma et al. (2006)*** but this would require the precise estimation of stimulus dependent correlations among the neurons.

We also did not find evidence for representing distributions via the DDC scheme during theta sequences. We note however, that on the behavioral time scale when averaging over many theta cycles, the sampling and DDC schemes become equivalent: in the sampling scheme the firing rate of neurons averaged across theta cycles is the expectation of their encoding basis functions under the encoded distribution. This way, sampling alternative trajectories in each theta cycle can be interpreted as DDC on the behavioral time scale with all computational advantages of this coding scheme (***Vértes and Sahani, 2019***). Similarly, sampling potential future trajectories naturally explains the emergence of successor representations on the behavioral time scale (***Stachenfeld et al., 2017***).

In standard sampling codes individual neurons correspond to variables and their activity (membrane potential or firing rate) represent the value of the represented variable which is very efficient for sampling from complex, high dimensional distributions (***Fiser et al., 2010; Orbán et al., 2016***). Here we take a population coding approach when individual neurons are associated with encoding basis functions and the population activity collectively encode the value of the variable (***Zemel et al., 1998; Savin and Deneve, 2014***). This scheme allows the hippocampal activity to efficiently encode the value of a low dimensional variable at high temporal precision.

**R**ecurrent networks can implement parallel chains of sampling from the posterior distribution of static (***Savin and Deneve, 2014***) or dynamic (***Kutschireiter et al., 2017***) generative models. Similar to the DDC scheme, these implementations would also encode uncertainty by the increase of the diversity of the co-active neurons. Thus, our data indicates that the hippocampus avoids sampling multiple, parallel chains for representing uncertainty in dynamical models and rather multiplexes sampling in time by collecting several samples subsequently at an accelerated temporal scale.

Our analysis leveraged the systematic increase in the uncertainty of the predicted states with time in dynamical models. The advantage of this approach is that we could analyse 1000s of theta cycles, much more than the typical number of trials in behavioral experiments where uncertainty is varied by manipulating stimulus parameters (e.g., image contrast; ***Orbán et al. 2016; Walker et al. 2020***). Uncertainty could also be manipulated during navigation by either introducing ambiguity regarding the spatial context (***Jezek et al., 2011***) or manipulating the volatility of the environment (***Miller et al., 2017***). These experiments would allow a more direct test of our theory by comparing changes in the neuronal activity with a behavioral readout of subjective uncertainty.

## Methods

### Theory

To study the neural signatures of the probabilistic coding schemes during hippocampal theta sequences we developed a coherent theoretical framework which assumes that the hippocampus implements a dynamical generative model of the environment. The animal uses this model to estimate its current spatial location and predict possible consequences of its future actions. Since multiple possible positions are consistent with recent sensory inputs and multiple options are available to choose from, representing these alternative possibilities, and their respective probabilities, in the neuronal activity is beneficial for efficient computations. Within this framework, we interpreted trajectories represented by the sequential population activity during theta oscillation as inferences and predictions in the dynamical generative model.

We define three hierarchical levels for this generative process (Fig. S1a). 1, We modeled the generation of *smooth planned trajectories* in the two-dimensional box, similar to the experimental setup, with a stochastic process. These trajectories represented the intended locations for the animal at discrete time steps. 2, We formulated the generation of *motion trajectories* via motor commands that are calculated as the difference between the planned trajectory and the position estimated from the sensory inputs. Again, this component was assumed to be stochastic due to noisy observations and motor commands. Calculating the correct motor command required the animal to update its position estimate at each time step and we assumed that the animal also maintained a representation of its inferred past and predicted future trajectory. 3, We modelled the generative process which *encodes the represented trajectories by neural activity*. Activity of a population of hippocampal neurons was assumed to be generated by either of the 4 different representational schemes as described below.

We implemented this generative model to synthesize both locomotion and neural data and used it to test contrasting predictions of different representational schemes. Importantly, the specific algorithm we used to synthesize motion trajectories and perform inference is not assumed to underlie the algorithmic steps implemented in the hippocampal network, it only provides sample trajectories from the distribution with the right summary statistics. The flexibility of this hierarchical framework enabled us to fit qualitatively the experimental data both at the behavioral (Fig. S2) and the neural (Fig. S3) level. In the following paragraphs we will describe these levels in detail.

### Generation of smooth planned trajectory

The planned trajectory was established at a temporal resolution corresponding to the length of a theta cycle, Δ.*t* = 0.1 s, and spanned a length *T* ≈ 1 s providing a planned position, 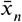 for any given theta cycle *n* (see Table 1 for a list of frequently used symbols). The planned trajectories were initialized from the center of a hypothetical 2 m×2 m box with a random initial velocity. Magnitude and direction of the velocity in subsequent steps were jointly sampled from their respective probability distributions. Specifically, at time step *n* we first updated the direction of motion by adding a random 2-dimensional vector of length 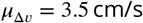 to the velocity 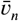. Next, we changed the speed, the magnitude of 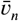, according to an Ornstein-Uhlenbeck process:

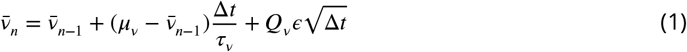

where 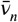 denotes the speed in theta cycle *n*. We used the parameters 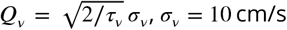 and *ϵ* ∼. 𝒩 (0, 1) when 2 cm/s ≤ ν ≤ 80 cm/s and *ϵ* = 0 otherwise. The planned trajectory was generated by discretized integration of the velocity signal:

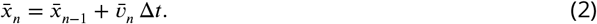

When the planned trajectory reached the boundary of the box, the trajectory was reflected from the walls by inverting the component of the velocity vector that was perpendicular to the wall.

**Table 1:**
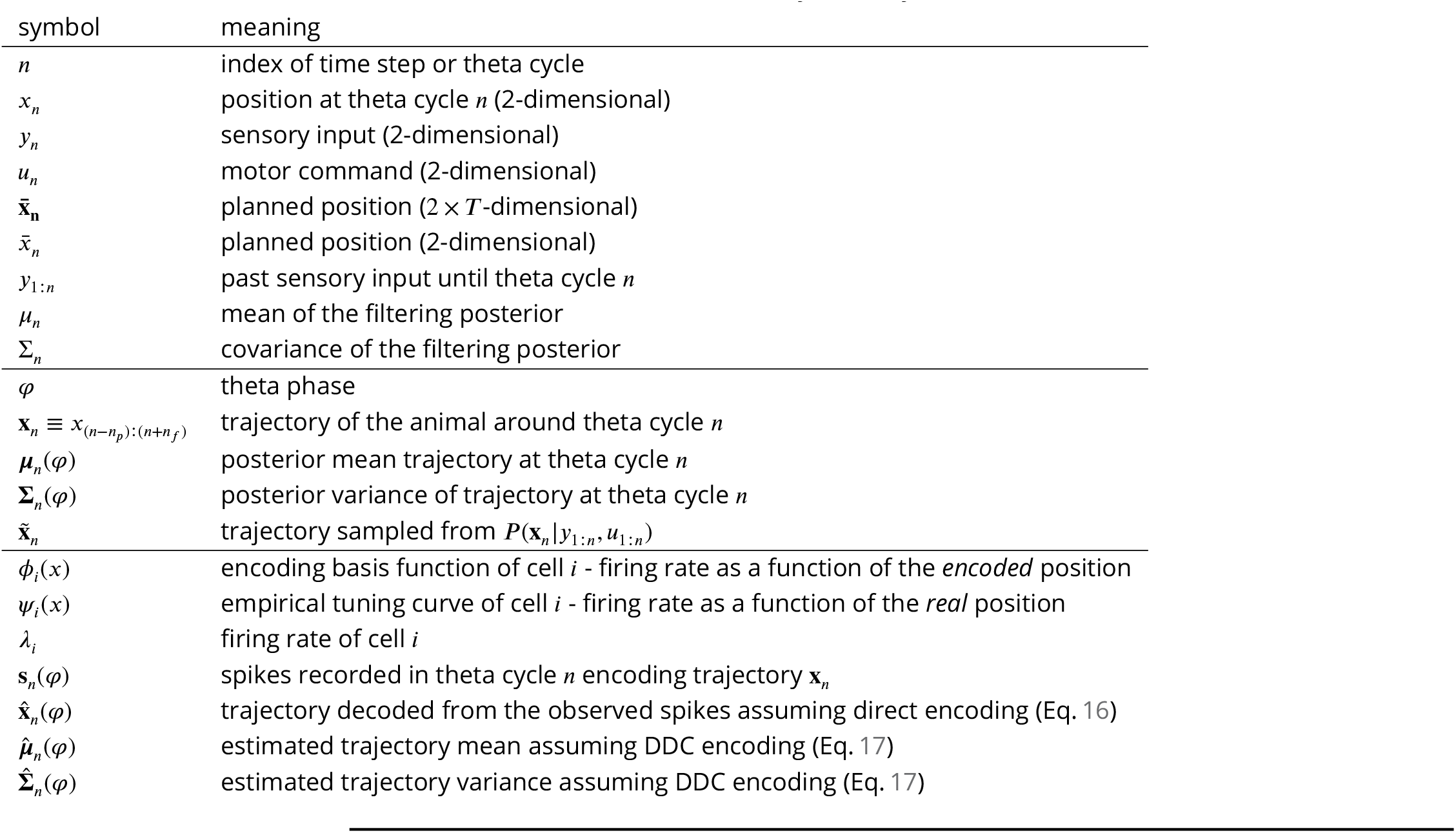
Summary of the symbols used in the model.

**Table 2:** P-values associated with Fig. 6. P-values for panels f,g and k were calculated using a one sample t-test. P-values for panel l were estimated by bootstrapping.

|  |  |  |  |  |  |  |  |
| --- | --- | --- | --- | --- | --- | --- | --- |
| panel: **f** |  |  |  |  |  |  |  |
| regular | jitter: 0 | 5 | 10 | 20 | 30 | 40 |  |
| 8.9e-10 | 1e-06 | 4.4e-07 | 1.5e-07 | 1e-05 | 5.7e-07 | 1.8e-07 |  |
| panel: **g** |  |  |  |  |  |  |  |
| regular | jitter: 0 | 5 | 10 | 20 | 30 | 40 |  |
| 0.0001 | 4e-15 | 3.9e-15 | 1e-13 | 3.4e-11 | 1.5e-11 | 2.4e-06 |  |
| panel: **k** |  |  |  |  |  |  |  |
| rat1 day1 | rat1 day2 | rat2 day1 | rat2 day2 | rat3 day1 | rat3 day2 | rat4 day1 | rat4 day2 |
| 5e-05 | 2.5e-18 | 2.5e-05 | 0.0001 | 2.1e-10 | 2.8e-07 | 1.4e-08 | 2.4e-06 |
| panel: **l** |  |  |  |  |  |  |  |
| rat1 day1 | rat1 day2 | rat2 day1 | rat2 day2 | rat3 day1 | rat3 day2 | rat4 day1 | rat4 day2 |
| <0.001 | <0.001 | <0.001 | <0.001 | <0.001 | <0.001 | <0.001 | <0.001 |

**Table 3:** P-values associated with Fig. S9. P-values for panels e,f and j were calculated using a one sample t-test. P-values for panel k were estimated by bootstrapping.

|  |  |  |  |  |  |  |  |
| --- | --- | --- | --- | --- | --- | --- | --- |
| panel: **e** |  |  |  |  |  |  |  |
| regular | jitter: 0 | 5 | 10 | 20 | 30 | 40 |  |
| 1.8e-05 | 1.4e-05 | 9e-05 | 4.8e-05 | 0.0016 | 0.0025 | 0.0006 |  |
| panel: **f** |  |  |  |  |  |  |  |
| regular | jitter: 0 | 5 | 10 | 20 | 30 | 40 |  |
| 0.01 | 4e-05 | 7e-05 | 0.0007 | 0.0006 | 0.014 | 0.16 |  |
| panel: **j** |  |  |  |  |  |  |  |
| rat1 day1 | rat1 day2 | rat2 day1 | rat2 day2 | rat3 day1 | rat3 day2 | rat4 day1 | rat4 day2 |
| 0.0018 | 4.5e-09 | 6e-08 | 4.9e-07 | 3.7e-19 | 4e-11 | 3.3e-25 | 2.4e-07 |
| panel: **k** |  |  |  |  |  |  |  |
| rat1 day1 | rat1 day2 | rat2 day1 | rat2 day2 | rat3 day1 | rat3 day2 | rat4 day1 | rat4 day2 |
| <0.001 | <0.001 | <0.001 | <0.001 | <0.001 | <0.001 | <0.001 | <0.001 |

Importantly, in any theta cycle multiple hypothetical planned trajectories could be generated by resampling the noise term, *ϵ* in Eq. 1. Moreover these planned trajectories can be elongated by recursively applying Eqs. 1-2. The planned trajectory influenced the motion of the simulated animal through the external control signal (motor command) as we describe it in the next paragraph.

### Generation of motion trajectories

We assumed that the simulated animal aims at following the planned trajectory but does not have access to its own location. Therefore the animal was assumed to infer the location from its noisy sensory inputs. To follow the planned trajectory, the simulated animal calculated motor commands to minimise the deviation between its planned and estimated locations.

To describe the transition between physical locations, *x*, we formulated a model where transitions were a result of motor commands, *u*_*n*_. For this, we adopted the standard linear Gaussian state space model:

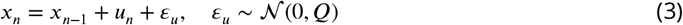

where *ε*_*u*_ represented noise coming from noisy execution of motor commands. The animal only had access to noisy sensory observations, *y*_*n*_ instead of locations:

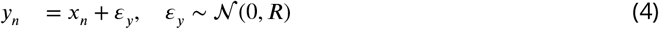

***Q*** and ***R*** are diagonal noise covariance matrices with ***Q***_*ii*_ = 2.25 cm^2^ and ***R***_*ii*_ = 225 cm^2^, indicating a more precise motor system.

Since location, *x*_*n*_, was not observed inference was required. This inference relied on estimates of the location in earlier theta cycles, the motor command, and the current sensory observation. The estimated location was represented by the Gaussian filtering posterior:

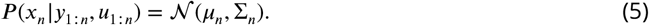

This posterior can be characterized by the mean estimated location *µ*_*n*_ and a covariance, Σ_*n*_, which is associated with the uncertainty of the estimate. These parameters were updated in each time step (theta cycle) using the standard Kalman filter algorithm (***Murphy, 2012***):

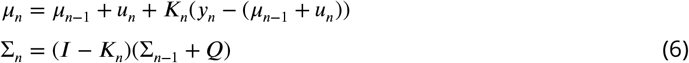

where *K*_*n*_ = (Σ_*n*−1_ + ***Q***)(Σ_*n*−1_ + ***Q*** + ***R***)^−1^ is the Kalman gain matrix.

The motor command, *u*_*n*_, was calculated by low-pass filtering the deviation between the planned position, 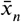, and the estimated position of the animal (posterior mean,*µ* _*n*−1_):

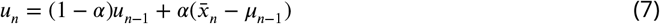

with *α* = 0.25. **R**elationship between the planned position 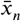, actual position *x*_*n*_, the sensory input *y*_*n*_, and the estimated location *P* (*x*_*n*_ I *y*_1:*n*_, *u*_1:*n*_) is depicted in Fig. S1b.

To make predictions about future positions, we defined the subjective trajectory of the animal, the distribution of trajectories consistent with all past observations and motor commands: *P* (**x**_*n*_ I *y*_1:*n*_, *u*_1:*n*_). This subjective trajectory is associated with a particular theta cycle: since it is estimated on a cycle-by-cycle manner we use the index *n* to distinguish trajectories at different cycles. Here 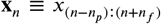 is a trajectory starting *n*_*p*_ steps in the past and ending *n*_*f*_ steps ahead in the future (Fig. S1c, Table 1). We call the distribution *P* (**x**_*n*_ I *y*_1:*n*_, *u*_1:*n*_) the *trajectory posterior*. We sampled trajectories from the posterior distribution by starting each trajectory from the posterior of current position (filtering posterior, Eq. 5) and proceeded first backward, sampling from the conditional smoothing posterior, and then forward, sampling from the generative model.

To sample the past component of the trajectory (*m < n*) we capitalised on the following relationship:

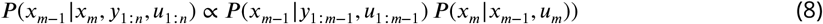

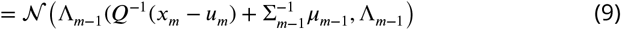

where the first factor of Eq. 8 is the filtering posterior (Eq. 5) and the second factor is defined by the generative process (Eq. 3) and 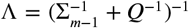. We started each trajectory by sampling its first point independently from the filtering posterior (Eq. 5) and applied Eq. 8 recursively to elongate the trajectory backward in time.

To generate samples in the forward direction (*m* ≥ *n*) we implemented an ancestral sampling approach. First, a hypothetical planned trajectory was generated as in Eq. 1-2 starting from the last planned location 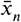. Next, we calculated the hypothetical future motor command, *u*_*m*+1_ based on the difference between the next planned location 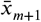 and current prediction for *m, x*_*m*_ as in Eq. 7. Finally, we sampled the next predicted position from the distribution

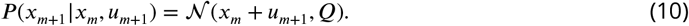

To elongate the trajectory further into the future we repeated this procedure multiple times.

We introduce ***µ***_*n*_(*φ*) = 𝔼 [*P* (**x**_*n*_ I*y*_1:*n*_, *u*_1:*n*_)] to denote the average over possible trajectories. *φ* indexes different parts of the trajectory and refers to the phase of the theta cycle at which the trajectory unofolds. Similarly, we also defined the covariance of the trajectories, **Σ**_*n*_(*φ*). We used an approximate, diagonal covariance matrix and ignored the covariances between different trajectories and theta phase. In our simulations we estimated ***µ***_*n*_(*φ*) and **Σ**_*n*_(*φ*) from 100 samples both for the past and for the future part of the represented trajectories.

The motion profile of the simulated animal, including the distribution and the auto-correlation of the speed, acceleration and heading was matched to empirical data from real animals (Fig. S2). Fig. S1c-d illustrates the inference process in the model by showing a short segment of the true trajectory of the simulated animal centered on its location at time step *n* as well as trajectories starting in the past and extending into the future sampled from the posterior distribution. As expected, the variance of these hypothetical trajectories **Σ**_*n*_(*φ*) increased consistently from the past towards the future (from the beginning to the end of the theta cycle; illustrated by the increasing diameter of the ellipses in Fig. S1d), while their mean ***µ***_*n*_(*φ*) tracked accurately the true trajectory of the animal (Fig. S1c-d). Mean trajectories and trajectories sampled from the trajectory posterior at subsequent theta cycles are compared in Fig. S1e-f and in Fig. 3a-b.

### Encoding the represented trajectory by the firing of place cells

We assumed that in each theta cycle the sequential activity of hippocampal place cells represents the temporally compressed trajectory posterior. The encoded trajectory was assumed to start in the past at the beginning of the theta cycle and arrive to the predicted future states (locations) by the end of the theta cycle (***Skaggs et al., 1996; Foster and Wilson, 2007***). Each model place cell *i* was associated with an encoding basis function, *ϕ*_*i*_(*x*), mapping the *encoded position* to the firing rate of cell *i*. Each basis function was generated as the sum of *K* Gaussian functions (subfields):

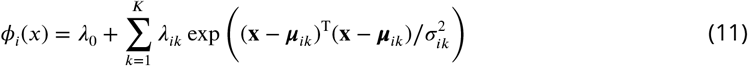

with the following choice of the parameters:

- The number of subfields *K* was sampled from a gamma distribution with parameters *α* = 0.57 and *β* = Ψ0.14 (***Rich et al., 2014***). We included only cells with at least one subfield within the arena (***K*** ≥ 1).
- The location of the subfields ***µ*** were distributed uniformly within the environment.
- The radius of each subfield, *σ*, was sampled uniformly between 10–30 cm.
- The maximum firing rate of each subfield *λ* was chosen uniformly on the range 5-15 Hz.
- The baseline firing rate *λ*_0_ was chosen uniformly on the range 0.1–0.25 Hz.

Examples of the encoding basis functions are shown in Fig. S7f (top row). Since the *encoded* positions can be different than the *measured* positions, the encoding basis function *ϕ*_*i*_(*x*) is not identical to the measured *tuning curve Ψ*_*i*_(*x*), which is defined as a mapping from the *measured* position to the firing rate. The exact relationship between tuning curves and the encoding basis functions depend on the way the estimated location is encoded in the population activity. Tuning curves estimated from the synthetic position data are compared with experimentally recorded place cell tuning curves in Fig. S3.

Importantly, in our model activity of place cells was determined by the inferred trajectories and not the motion trajectory of the animal. The way the trajectories were encoded by the activity of place cells was different in the four encoding schemes:

1. In the *mean* encoding scheme the instantaneous firing rate of the place cells was determined by the mean of the trajectory posterior

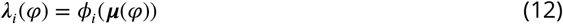

That is, the cell’s firing rate changed within the theta cycle according to the value of its basis function at the different points of the mean inferred trajectory.
2. In the *product* scheme (***Ma et al., 2006***) the firing rate was controlled both by the posterior mean and variance of the trajectory:

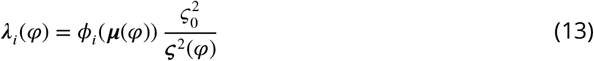

where 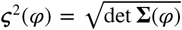 and *ς*_0_ = 5 cm. This is similar to the mean encoding model, except that the population firing rate is scaled by the inverse of the posterior variance.
3. In the *DDC* scheme (***Zemel et al., 1998; Vértes and Sahani, 2018***) the instantaneous firing rate of cell *i* is the expectation of the basis function *i* under the trajectory posterior at the encoded time point (that is, the overlap between the basis function and the posterior):

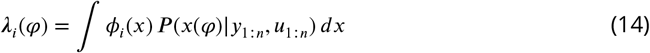
4. In the *sampling* scheme, the encoded trajectory was sampled from the trajectory posterior, 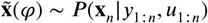 and the instantaneous firing rate was the function of the sampled trajectory:

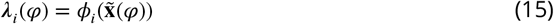

In each encoding model, spikes were generated from the instantantous firing rate ***λ***_*i*_(*φ*) as an inhomogeneous Poisson process:

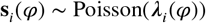

### Decoding

To discriminate encoding schemes from each other we decoded position information both from experimentally recorded and from synthesized hippocampal neuronal activity. We performed decoding based on two different assumptions: First, assuming that a single position is encoded by the population activity (consistent with the mean and sampling schemes). Second, assuming that a distribution is encoded via the DDC scheme. We used static decoders to ensure that the variance of the decoder is independent of the theta phase as opposed to dynamic decoders, where the variance can be larger around the boundaries of the decoded segments.

### Single point decoding

We performed static Bayesian decoding independently in each temporal bin at different phases of the theta cycle. The estimated position at theta phase *φ* is the mean of the posterior distribution calculated using Bayes rule:

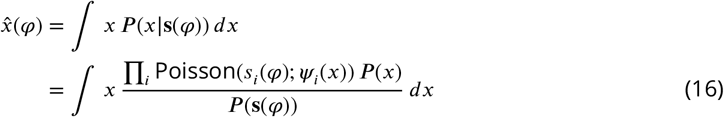

where the prior *P* (*x*) was the estimated occupancy map and we used a Poisson likelihood with spatial tuning curves *Ψ*_*i*_(*x*), estimated from the smoothed (10 cm Gaussian kernel) and binned spike counts. We binned spikes either into windows of fixed duration (20 ms, Fig. 2a-c) fixed theta phase (120^°^, Fig. 3h and Fig. 4) or into three bins with equal number of spike counts within a theta cycle (Fig. 5-6 and everywhere else). When calculating the spread of the decoded locations (Fig. 2g, Fig. 3h), we controlled for the possible biases introduced by theta phase modulation of the firing rates by randomly downsampling the data to match the spike count histograms across theta phase.

### DDC decoding

The DDC decoder assumes that at each time point in the theta cycle an isotropic Gaussian distribution over the locations is encoded by the population activity via Eq. 14 and we aim at recovering the mean and the variance of the encoded distribution. To ensure that the theta phase dependence of the firing rates does not introduce a bias in the decoded variance, we divided each theta cycle into three windows (early, middle and late) with equal number of spikes. As linear decoding of DDC codes from spikes (***Vértes and Sahani, 2018***) is very inaccurate, we performed maximum likelihood decoding of the parameters of the encoded distribution. The estimated mean and the variance in bin *φ* is:

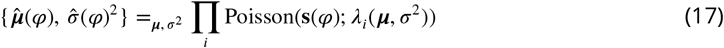

where

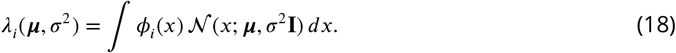

Here, *ϕ*_*i*_(*x*) is the basis function of neuron *i* used in the encoding process (Eq. 14). We numerically searched for the maximum likelihood parameters with constraints σ ∈ (0, 200) (cm) and *σ* E (3, 200) (cm) using quasi-Newton optimizer and a finite-difference approximation for the gradients.

In practice we do not have access to the encoding basis functions, *ϕ*_*i*_(*x*), only to the empirical tuning curves, *Ψ* (*x*). Based on the synthetic data we found that the tuning curves are typically more dispersed than the basis functions when the encoded and the measured location is not identical or when the encoded distributions have nonzero variance (Fig. S7e, g, Supplemental Note). The difference between the size of the basis functions used for encoding and decoding introduces a bias in decoding the variance of the distribution (Fig. S7a, c). To reduce this bias, we devised a non-parametric deconvolution algorithm that could estimate the encoding basis functions from the empirical tuning curves (Fig. S7d-f, Supplemental Note). Although we demonstrated on synthetic data that the bias of the decoder can be eliminated by using these estimated basis functions (Fig. S7d), we obtained similar results with either the estimated basis functions or the empirical tuning curves. Therefore throughout the paper we show DDC decoding results obtained using the empirical tuning curves and show decoding with the estimated basis functions in the Supplemental material (Fig. S8, Supplemental Note)

### Analysis of trajectory variability

To discriminate the mean scheme from the sampling scheme we introduced the excess variability index (EV-index). The EV-index is the difference between the cycle-to-cycle variability and the trajectory encoding error and is positive for sampling and negative for the mean scheme. In this section we provide definitions for these quantities, derive their expected value for the sampling and the mean scheme and show how the EV-index can be estimated from data.

#### Cycle-to-cycle variability (*CCV*)

We defined cycle-to-cycle variability (*χ*) as the difference between the trajectories decoded from the neural activity in two subsequent theta cycles (Fig. S1f):

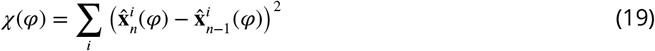

where *φ* is the theta phase and the index *i* runs over the 2 dimensions. As we show in Supplemental Note, for the mean encoding scheme the expected value of the cycle-to-cycle variability is the sum of two terms:

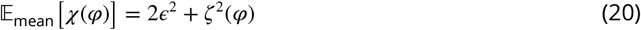

where 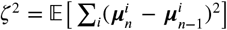 is the expected change of the encoded mean trajectory between subsequent theta cycles, *ϵ*^2^ is the error of the decoder reflecting the finite number of observed neurons in the population and their stochastic spiking. In the above equations the expectation runs across theta cycles. Since each theta cycle is divided into equal spike count bins, the decoder variance is independent of the theta phase *φ*. Conversely, ς^2^ increases with *φ* (Fig. S1g) as new observations have larger effect on uncertain future predictions than on the estimated past positions.

Although our derivations use the expected value (mean across theta cycles), in practice we often found that the median is more robust to outliers and thus in Fig. 6 and Fig. S9 we also show results with median instead of mean.

When estimating the cycle-to-cycle variability for the sampling scheme, where the population activity encodes independent samples drawn from the trajectory posterior, an additional term appears:

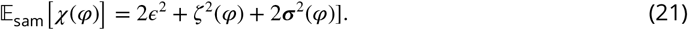

Here ***σ***^2^(*φ*) = 𝔼[∑_*i*_ **Σ**^*i*^(*φ*)] is the total posterior variance, which reflects the variance coming from the uncertainty of the inference.

Independent estimation of the variance in the encoded trajectory is challenging but we can exploit insights obtained from synthetic data. In our synthetic dataset we found that the trajectory change is proportional to the posterior variance:

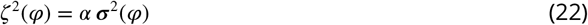

with the proportionality constant 0 *< α <* 1 (Fig. S1g). Using this insight, we can simplify our treatment: by substituting Eq. 22 into Eqs 20-21, we can see that in both coding schemes the cycle-to-cycle variability increases with the theta phase, and the magnitude of this increase, proportional to the total posterior variance ***σ***^2^(*φ*), can discriminate the two coding schemes. In order to obtain an independent estimate of ***σ***^2^(*φ*) we introduce another measure, the trajectory encoding error.

#### Trajectory encoding error (*TEE*)

We defined trajectory encoding error (*γ*) as the expected difference between the true 2-dimensional trajectory of the rat, **x**_*n*_, and the trajectory decoded from the neural activity 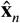 (Fig. S1e):

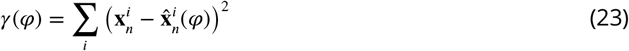

When comparing decoded and physical trajectories, we assumed a fixed correspondence between theta phase *φ* and temporal shift along the true trajectory. Specifically, for each animal we first calculated the average decoding error for early, mid and late phase spikes with respect to the true position temporally shifted along the motion trajectory (Fig. S4). Next, we determined the temporal shift Δ*t* that minimised the decoding error separately for early, mid and late theta phases and used this Δ.*t* to calculate *γ*.

In the case of mean encoding the trajectory encoding error is the sum of two terms:

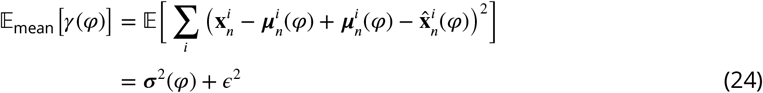

where we used the fact that the encoded trajectory is the mean ***µ***_*n*_(*φ*), and if the model of the animal is consistent, then the expected difference between the posterior mean and the true location equals the variance of the posterior, 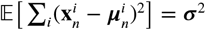.

In the case of sampling the encoded trajectory is 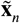 and the trajectory encoding error is increased by the difference between the mean and the sampled trajectory:

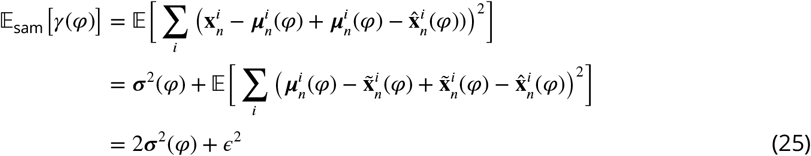

### Decoding error

To compare cycle-to-cycle variability with trajectory encoding error we have to estimate the decoding error, *ϵ*^2^. A lower bound to the decoding error is given by the Cramér-**R**ao bound (***Dayan and Abbott, 2001***), but in our simulations the actual decoding error was slightly larger than this bound. Underestimating the decoding error would bias the excess variability towards more positive values (see below), which we wanted to avoid as it would provide false evidence for the sampling scheme.

To obtain a reasonable upper bound instead, we note, that both *χ* and *γ* were evaluated in three different phases of the theta cycles: early (*φ*_1_), mid (*φ*_2_) and late (*φ*_3_). At early theta phases when encoding past positions the posterior variance ***σ***^2^ is small and thus both *χ* and *γ* are dominated by *ϵ*^2^. Thus, we estimated the decoding error from the measured cycle-to-cycle variability and trajectory encoding error at early theta phase.

Furthermore, to compare cycle-to-cycle variability with trajectory encoding error we defined the compensated cycle-to-cycle variability and trajectory encoding error by subtracting the estimated decoding error:

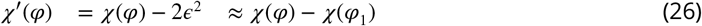

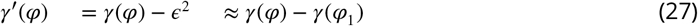

#### Excess variability

We define the excess variability as the difference between *ϵ′*^’^ and *γ′*^’^:

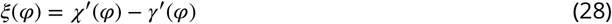

Substituting Eq. 20-21, Eq. 24-25 and Eq. 26-27 to Eq. 28 we can obtain the expectation of the excess variability in the sampling and the mean encoding scheme:

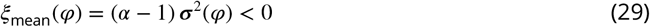

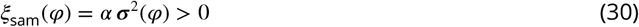

The excess variability is positive for sampling and negative for mean encoding. As ***σ***^2^ is expected to increase within a theta cycle, the excess variability is most distinctive at late theta phases. Therefore, troughout the paper we defined the *EV-index* as the excess variability at late theta phases:

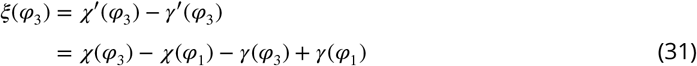

Importantly, all terms in Eq. 31 can be measured directly from the data.

We calculated the EV-index either using all theta cycles (Fig. S9) or using theta cycles with high spike counts in order to mitigate the effect of the decoding error (Fig. 6). To reduce the effect of outliers, we also reported the median of the EV-index in Figures 6l and S9k. Error bars on the EV-index (Fig. 6f-g, k, Fig. S9e-f, j) reflect the standard error across the theta cycles. When showing the median across the theta cycles (Fig. 6l, Fig. S9k), the error bars indicate the 5% and 95% confidence intervals estimated by bootstrapping. Specifically, we obtained 1000 pseudo-datasets by discarding randomly selected half of the theta cycles and calculating the EV-index of the remaining data.

## Data analysis

### Processing experimental data

To test the predictions of the theory, we analysed the dataset recorded by ***Pfeiffer and Foster*** (***2013***). In short, rats were required to collect food reward from one of the 36 uniformly distributed food wells alternating between random forging and spatial memory task. The rat’s position and head direction were determined via two distinctly coloured, head-mounted LEDs recorded by an overhead video camera and digitized at 30 Hz. Neural activity was recorded by 40 independently adjustable tetrodes targeting the left and the right hippocampi. Local field potential (LFP) was recorded on one representative electrode, digitally filtered between and 500 Hz and recorded at 3,255 Hz. Individual units were identified by manual clustering based on spike waveform peak amplitudes based on the signals digitalized at 32,556 Hz as in ***Pfeiffer and Foster*** (***2013***).

The raw position signal was filtered with a 250 ms Gaussian kernel and instantaneous speed and motion direction was calculated from the smooth position signal. We restricted the analysis to run periods with υ *>t* 5 cm/s for at least 1 s and stops with duration of *δ >* 0.2 s between two subsequent run epochs. In this study we included only putative excitatory neurons (on the basis of spike width and mean firing rate (***Pfeiffer and Foster, 2013***)) with at least 200 spikes during the analysed run epochs. Position was binned (5 cm) and spatial occupancy map was calculated as the smoothed (10 cm Gaussian kernel) histogram of the time spent in each bin. Position tuning curves (ratemaps, *Ψ* (*x*)) were calculated as the smoothed (10 cm Gaussian kernel) histogram of firing activity normalized by the occupancy map and we used *Ψ*_min_ = 0.1 Hz wherever *Ψ* (*x*) *<* 0.1 Hz.

Theta phase was calculated by applying Hilbert transformation on band pass filtered (4–12 Hz) LFP signal. To find the starting phase of the theta sequences in each animal, we calculated the forward bias by decoding spikes using 120^°^windows advanced in 45^°^increments (Fig. 4h). The forward (lateral) bias of the decoder was defined as the average of the error between the decoded and the actual position of the animal parallel (perpendicular) to the motion direction of the animal. Theta start was defined as 135^°^after the peak of the cosine function (Fig. 4f) fitted to the forward bias. We used two different strategies to avoid biases related to systemic changes in the firing rate during decoding: (1) We subsampled the data in order to match the distribution of spike counts across theta phases. (2) When analysing early, mid and late theta phases, we divided each theta cycle into three periods of equal spike count. The decoding spread was defined as det(Σ)^¼^ where Σ is the covariance matrix of the decoded positions.

To calculate the theta phase dependence of the place field size (Fig. 2e) we estimated the place fields in the three theta phase (early, mid and late) using the position of the rat shifted in time with Δ*t*(*φ*) that minimized the median error between the decoded and temporally shifted position in that session (Fig. 2d). Similarly, when we calculated trajectory encoding error, we compared the decoded position in each phase bin to the real position shifted in time with the same Δ*t*(*φ*).

### Processing synthetic data

We simulated the movement of the animal using Eq. 1-7 in a 2×2 m open arena, except for Fig. 3d-e, where we used a 2 ×0.1 m long linear track. We used the same procedure to analyse the synthetic data as we applied to the experimental data: we filtered the raw position signal, calculated the speed and motion direction and estimated the spatial occupancy maps and and position tuning curves from the generated spikes. Importantly, we used these estimated tuning curves and not the true, noiseless tuning functions for decoding position from spikes in the synthetic data. In our model each theta cycle was 100 ms long (but also see an alternative variant below) and the encoded trajectory spanned 2 s centered on the current position of the animal.

To test the robustness of the EV-index to variations across theta cycles, we added variability to the theta cycles in two different ways: First, the duration and the content of each theta cycle was varied stochastically. Specifically, we varied the duration of each theta cycle (80–160 ms), the total length of the encoded trajectory (1–3 s) with constraining the encoded trajectory to start in the past and end in the future. Second, a uniformly distributed random jitter (0–40 ms) was added to the true time of the theta cycle boundaries before binning the spikes according to their theta phase.

## Acknowledgements

We thank Brad E Pfeiffer and David J Foster for kindly sharing their data and Andres D Grosmark and György Buzsáki for making their data publicly available. We thank Márton Kis and Judit K Makara for useful discussions; Mihály Bányai and Judit K Makara for comments on a previous version of the manuscript. This work was supported by an NKFIH fellowship (PD-125386, FK-125324; BBU), by the Hungarian Brain **R**esearch Program (2017-1.2.1-NKP-2017-00002, GO and KTIA-NAP-12-2-201, BBU and GO).

## Author contributions

BBU and GO S.P. conceived and designed the study. BBU developed the model and performed the analysis with input from GO. BBU and GO wrote the paper.

## Supplemental Material

### Supplemental Note S1 – Product coding scheme

In a product representation, a probability distribution over a feature, such as the position *x*, is encoded by the product of the encoding basis functions, *ϕ*_*i*_(*x*):

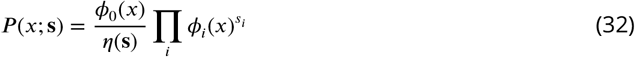

where *s*_*i*_ is the number of spikes fired by neuron *i* and 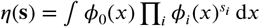 is the normalizing factor. Alternatively, by defining a set of sufficient statistics, *h*_*i*_(*x*) = log *ϕ*_*i*_(*x*), the same distribution can be expressed in the standard, exponential form:

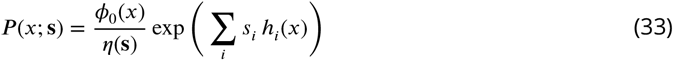

where the neuronal spikes serve as the canonical or exponential parameters of the distribution (***Wainwright and Jordan, 2008***). Intuitively, increasing the number of spikes **s** in the population adds more and more terms in Eqs.32-33 leading to an increase in the sharpness of the represented distribution. In the special case where the tuning functions are Gaussians with variances 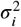 and the likelihood is conditionally independent Poisson, the represented distribution is also Gaussian with a variance, σ^2^(**s**) that is inversely proportional to the total spike count (***Ma et al., 2006***):

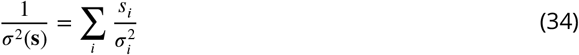

Here we show numerically that this relationship still holds for more general, multimodal basis functions, similar to the empirical tuning curves found in our dataset (Fig. 2b). We used 110 tuning curves from an example session with mean firing rate 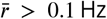 and tuning curve peak *r*_max_ *>* 10 Hz (Fig. S6a), we normalised the tuning curves and used their logarithm as sufficient statistics to encode distributions in the form of Eq. 33:

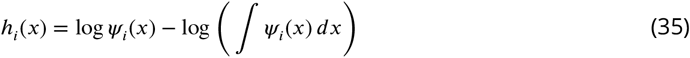

We used isotropic 2-dimensional Gaussians with a wide range of mean (20 cm ≥ ≥ 180 cm) and variance (4 cm^2^ ≥ *σ*^2^ ≥ 1600 cm^2^) as target distributions and encoded them in the spiking activity of the neurons. Eq. 33 defines how a set of observed spikes (canonical parameters) can be inteprreted as encoding a probability distribution. However, this mapping is not invertible and finding a set of spikes approximating an arbitrary target distribution is computationally challenging. We solved this problem by iteratively and greedily selecting spikes to minimise the Kullback-Leibler divergence between the encoded distribution, ***Q***(*x*) ∝ exp (∑_*i*_ *s*_*i*_ *h*_*i*_(*x*)), and the Gaussian target distribution, *P* (*x*). Specifically, we first selected a single neuron whose spike minimised D_KL_(***Q***, *P*), then added more spikes one by one until the additional spikes stopped decreasing D_KL_(***Q***, *P*).

We found that the approximations were reasonably accurate, especially for distributions with smaller variance (Fig. S6b). Importantly, the average number of spikes used for encoding decreased systematically with the variance of the encoded distribution (Fig. S6c). Moreover, we found that even for highly variable basis functions, the total spike count in the population was well approximated as a linear function of the target precision, as expected from the theory for uniform basis functions (Fig. S6d; Eq. 34). Therefore we used the systematic variation of the total spike count in the population (population gain) with uncertainty as a hallmark of the product scheme.

When encoding trajectories in the population activity we used a simplified encoding model and scaled the gain of the neurons by the instantaneous variance of the encoded distribution (Eq. 13).

### Supplemental Note S2 – DDC coding scheme

In the DDC coding scheme the firing rate of neuron *i* corresponds to the expectation of its basis function under the encoded distribution (***Zemel et al., 1998; Vértes and Sahani, 2018***):

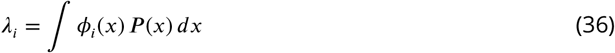

This encoding scheme has complementary properties to the product scheme, as it represents exponential family distributions using their mean parameters instead of their canonical, natural parameters (***Wainwright and Jordan, 2008***). Eq. 36 defines a mapping from a distribution to firing rates, but the reverse mapping, from the rates to the encoded distribution is not trivial. When the firing rates are observed then the expectation of any nonlinear function *f* (·) under the encoded distribution, including its moments, the mean and the covariance, can be calculated as a linear combination of the rates (***Vértes and Sahani, 2018***):

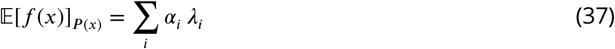

where *f* (*x*) ≈ ∑ _*i*_ *α*_*i*_ *ϕ*_*i*_(*x*). However, in the experimental data only the discrete spike counts are observed and the underlying firing rates are hidden. In this case Eq. 37 can be computed as an expectation under the posterior of the firing rates *P* (***λ***I**s**):

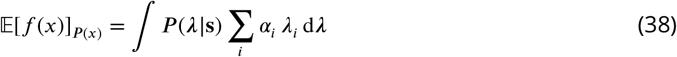

with

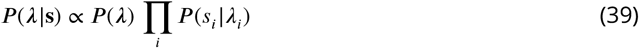

Here the prior over the firing rates *P* (***λ***) (i.e., the coactivity of neurons with different tuning curves or place fields) introduces strong correlations in the posterior and thus using a factorised approximation of Eq. 38 leads to a substantial loss of information (***Ujfalussy et al., 2015***). Therefore, instead of following this Bayesian approach, we aimed at directly estimating the parameters of the distribution encoded by the spikes observed in a neuronal population using a maximum likelihood decoder (Eq. 17).

We tested the performance of the maximum likelihood DDC decoder by encoding isotropic 2-dimensional Gaussian distributions with a wide range of mean (20 cm ≥ µ ≥ 180 cm) and variance (4 cm^2^ ≥ *σ*^2^ ≥ 400 cm^2^) as target in the spiking activity of a neural population using 200 encoding basis functions similar to the empirical tuning curves recorded experimentally (Fig. S7g, top). We calculated the firing rate of the neurons according to Eq. 36 and generated Poisson spike counts in Δ*t* = 20 − 100 ms time bins. We found that on average, we could accurately estimate the encoded mean and SD of the target distributions when we used the same basis functions during encoding and decoding at least for Δ*t* ≥ 50 ms. (Fig. S7b). However, in practice, we do not have access to the encoding basis functions, only the empirical tuning curves, substantially wider than the basis functions (Fig. S7c). When we used the wider tuning curves for decoding, the estimate of the SD became substantially biased: the decoded SD could be on average 5-7 cm below the target SD even for Δ*t* = 100 ms (Fig. S7c).

To identify DDC scheme in the data it is important to be able to reduce this decoding bias and accurately estimate the SD of the encoded distribution even in small time windows. We speculated that this bias is due to the difference between the wider empirical tuning curves used for decoding and the narrower basis functions used for encoding the target distributions. This effect is caused by encoding locations other than the actual position of the animal and by encoding distributions with non-zero variance using the DDC scheme. The decoder tries to match the overlap between the tuning functions and the estimated distribution with the observed spike counts. When the tuning functions used for DDC-decoding are wider than the encoding tuning functions, the decoder will compensate this by choosing a narrower decoded distribution with a similar overlap with the decoding tuning functions (Fig. S7a). Thus, the decoder will be biased towards lower variances, leading to systematic underestimation of the variance of the encoded distribution.

In order to reduce this bias we aimed at estimating the true encoding basis functions from the recorded neuronal activity. Note, that deriving an optimal estimator is hindered by the fact that in Eq. 36 neither the tuning functions nor the encoded distributions or the true firing rates are observed. Therefore we developed an approximate algorithm to sharpen the empirical tuning curves and reconstruct the original tuning functions. The aim was to find a flexible algorithm that preserves the relative mass of different modes of the tuning curves but sharpens all of them to a similar degree. Our method was inspired by particle filtering and the use of annealing methods in sampling (***Murray, 2007***): we first generated a set of samples from the empirical tuning curves, and then added random noise to propagated the samples towards the desired distribution:

1. We generated *N* = 10, 000 samples from the empirical tuning curve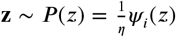, where *η*= ∫ *Ψ*_*i*_(*z*)d*z* is the normalization constant to convert the tuning curve to a distribution.
2. We added a Gaussian random noise to the samples **z**^′^ = **z** + *ϵ* with *ϵ*∼ 𝒩 (0, *σ*^2^) where the proposal variance, *σ*^2^ controls the degree of mixing between the different modes. We used *σ* = 10 cm.
3. We accepted the new sample 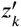 with probability 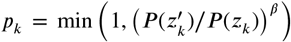 where the parameter *β* controls the direction and degree of sharpening. We used *β* = 3.
4. We repeated steps 2. and 3. for a number of iterations, not necessarily until convergence.
5. Finally, we defined the estimated basis functions as the smoothed (5 cm Gaussian kernel) histogram of the number of samples in 5 cm spatial bins.

Fig. S7g shows the tuning functions of a few example cells from our synthetic dataset (top) together with the measured tuning curves (middle) and the result of the sharpening algorithm after 3 and 6 steps (bottom). We evaluated the sharpening algorithm by comparing the relative size of the tuning curves and the tuning functions and the average difference between them. We found that 6 iterations of the algorithm eliminates overestimation bias of the tuning curve size (Fig. S7e) and 3-6 iteration minimized the estimation error (Fig. S7f). Using the 3-step sharpened tuning curves also improved the performance of the DDC-decoder as it substantially reduced the bias when decoding the SD of the distribution (Fig. S7d). Therefore we repeated the DDC decoding analysis shown in Fig. 5 using the 3-step sharpened tuning curves in Fig. S8.

Even with this correction, trial by trial decoding of the represented distributions was not possible on short time scales. However, when averaged across 1000s of theta cycles, the decoder became sufficiently sensitive to the encoded SD to identify systematic changes in the represented uncertainty. Therefore, we used the average SD decoded from the neural activity as a hallmark of DDC encoding.

### Supplemental Note S3 – Background for the calculation of EV-index

#### Cycle-to-cycle variability

Here we derive the expectation of Eq. 19 in the case of mean encoding scheme, when the population activity encodes the posterior mean trajectory ***µ***_*n*_:

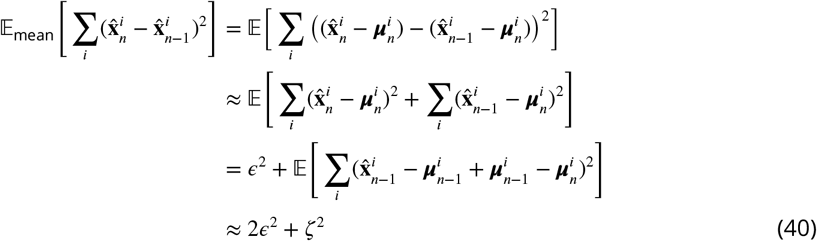

where 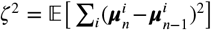 is the expected change of the posterior mean trajectory due to sensory inputs, and we omitted second order terms including the correlations between the decoding errors in subsequent theta cycles and the correlation between decoding error and trajectory change.

The same expectation in the case of sampling, when the encoded location 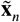 is sampled from the posterior:

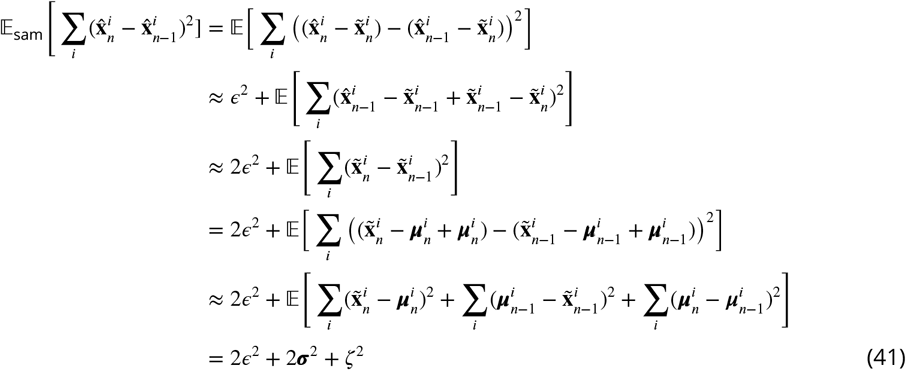

where ***σ***^2^ = 𝔼 [∑*i* **Σ**^*i*^ measures the total variance of the posterior and we omitted all second order terms. We found that the contribution of second order terms are small in most cases, except for the *border-effect* which captures the correlation of the decoding error in two subsequent theta cycles. The relatively strong positive correlation is explained by the bias of the decoder near the borders of the arena. Importantly, the contribution of this term is independent of theta phase and thus does not influence our analysis. We also assumed that the sample autocorrelation is near zero i.e., the samples drawn in subsequent theta cycles are independent.

## Supplemental Figures

**Figure S1:**
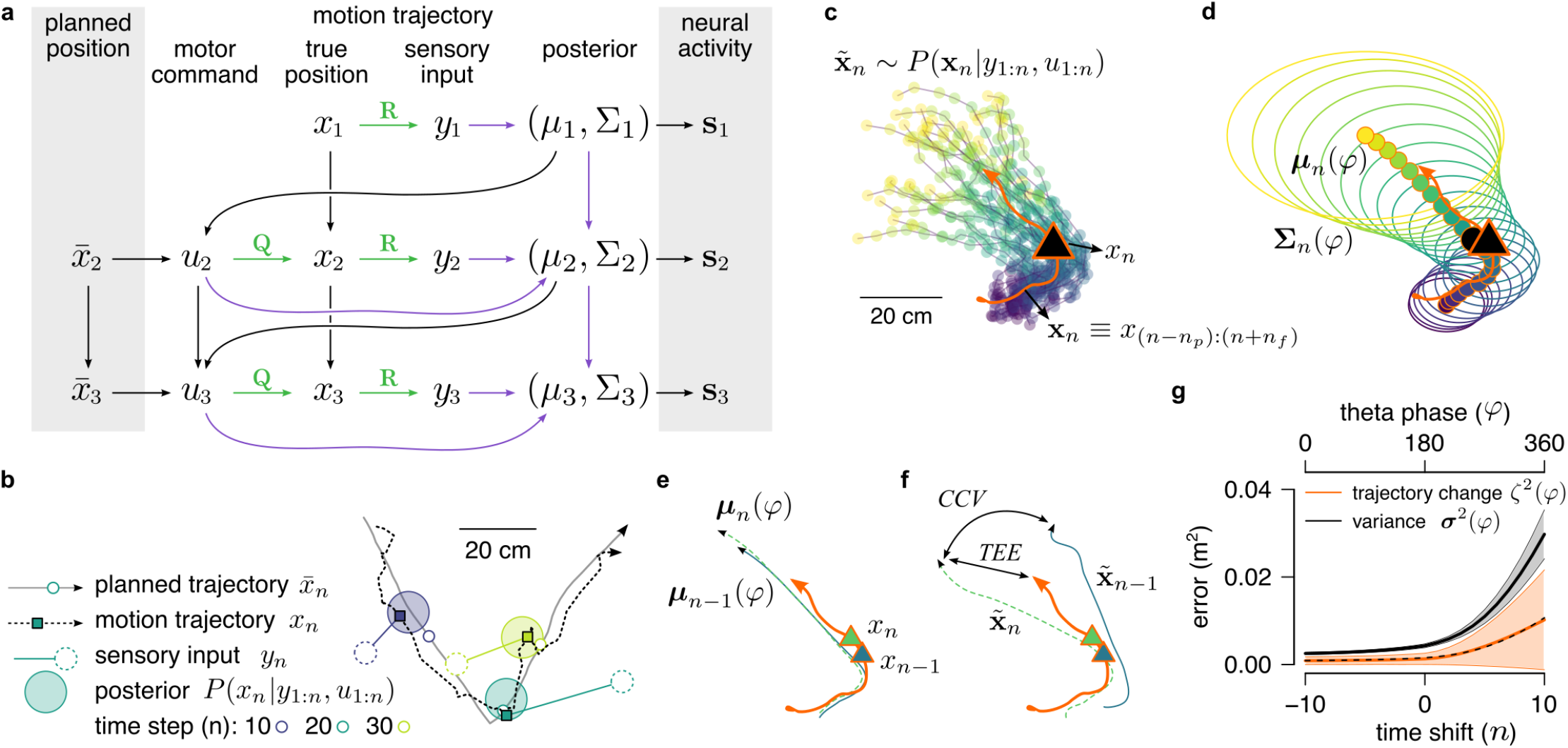
Inference and movement in the generative model. **a** Graphical model of the processes underlying the generation of the simulated animal’s trajectory. Atthe unknown true location, *x*_*t*_, the animal receives a noisy sensory input, *y*_*t*_, and combines it with the previous motor command, *u*_*t*_, to update its current estimate of its location *P(x*_*t*_ |*y*_*l*_._*t*_,*u*_*l*_._*t*_*) = 𝒩 (µ*_*t*_,Σ*t*). Based on its current position estimate and the intended position for the next time step, 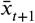 it calculates the next motor command, *u*_*t+l*_, that is used to generate the new position *x*_*r*+1_. Green arrows represent sensory and motor noise. Purple arrows represent inference of *P(x*_*t*_ |*y*_1_._*t*_,*u*_1_._*t*_). Neuronal activity **s**_*r*_ depends on the estimated position. **b** A 4s segment of planned trajectory (grey) and motion trajectory (dashed) together with the true, planned and inferred positions and the sensory input at 3 different time points separated by 1 s (colors). **c** The motion trajectory of the simulated animal (orange) with its inferred past (blue) and predicted future (yellow) positions (colors) represented as 25 trajectories sampled at position *x*_*t*_ (triangle). **d** Same as B except that distribution of trajectories are represented by covariance ellipses. **e** True trajectory (orange) and posterior mean trajectories at two subsequent timesteps (theta cycles, blue and green). **f** Similar to D but with trajectories sampled independently from the corresponding posterior distribution. Trajectory encoding error(*TFF*) denotes the difference between the true and the sampled trajectory whereas cycle-to-cycle variability (*CCU*) is the difference between trajectories sampled in subsequent theta cycles. **g** Posterior variance ***σ***^2^ and change in the mean trajectory *ζ*^2^ as a function of the time shift (sensory inputs and motor commands are observed till *n =* 0). Shaded region shows SD. Trajectories with time shift −10 ≤ *n* ≤ 10 (corresponding to −1≤ Δ *t* ≤ 1 s) are encoded in each theta cycle between 0^°^≤ *φ≤*360^°^. Dashed line shows the linear fit *ζ*^2^ ≈ *ασ*^*2*^.

**Figure S2:**
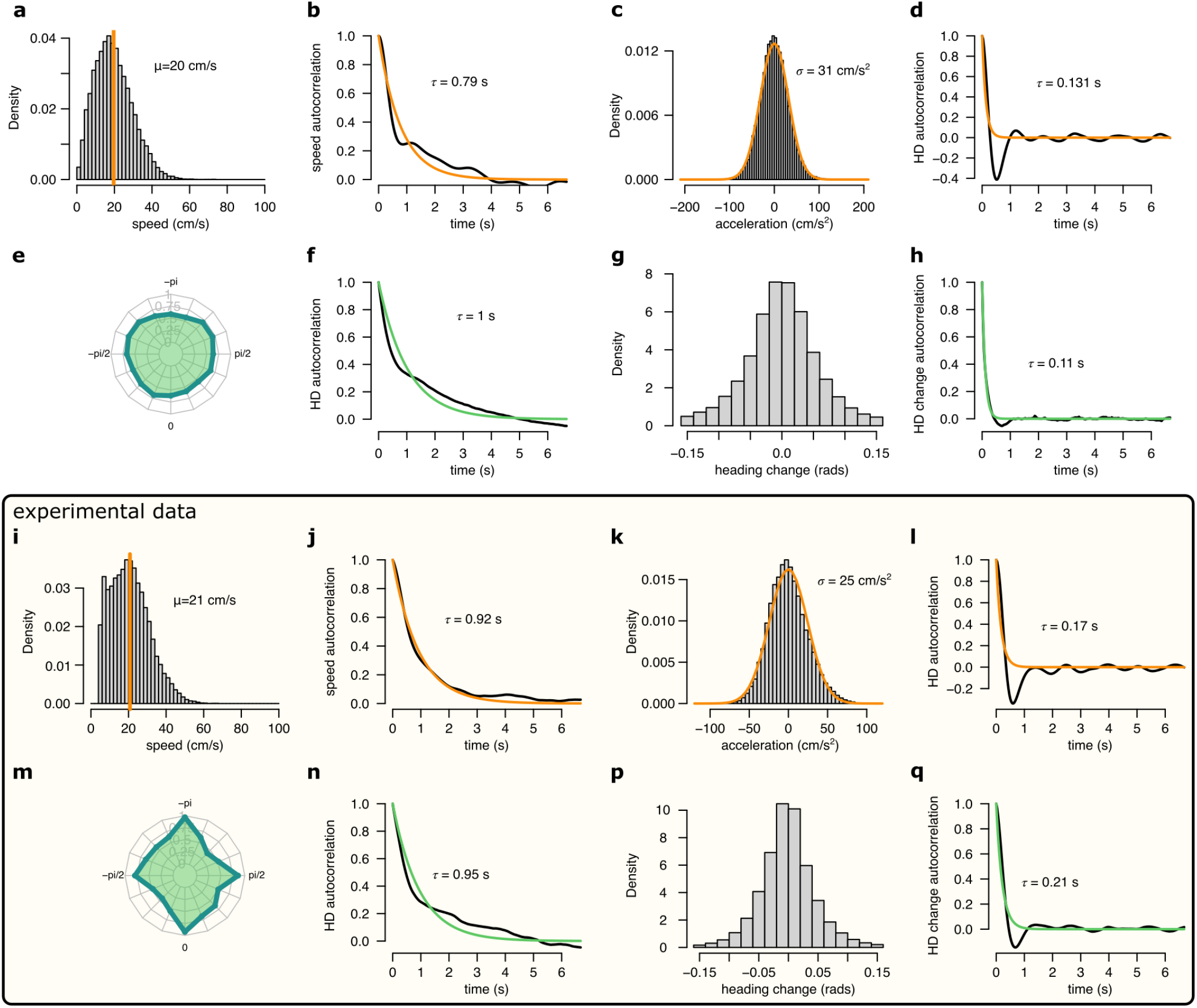
Comparison of the motion profile of the simulated animal and one of the analysed experimental sessions. **a-h** Motion profile of the simulated animal. **a** Histogram of the running speed. **b** Autocorrelation of the running speed. **c** Histogram of the acceleration. **d** Autocorrelation of the acceleration. **e** Polar histogram of the running directions. **f** Autocorrelation of the head direction. **g** Histogram of the head direction change per time step (1/30s). **h** Autocorrelation of the head direction change. **i-q** Same as a-h for experimental session rat 1 day 2. Note, that for the experimental data we only analysed continuous running periods, where *υ >* 5 cm/s.

**Figure S3:**
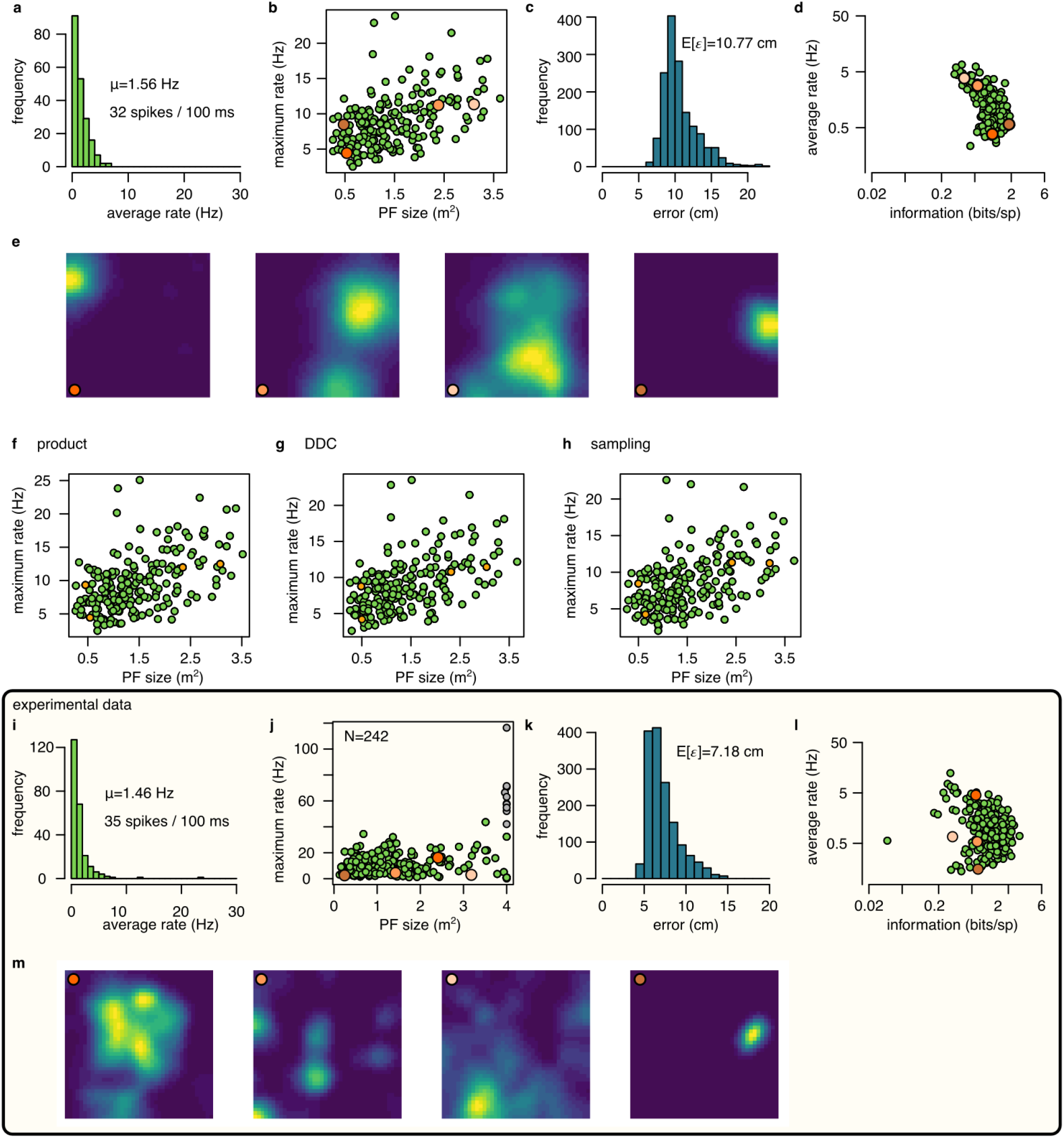
Place cell firing in the synthetic and in the experimental data. **a-e** Place cell activity in the simulated data using mean encoding. **a** Histogram of the average firing rate of the 200 simulated neurons. Numbers indicate the average firing rate across cells and the average number of spikes in a 100 ms theta cycle. **b** Maximum firing rate as a function of place field size (area over the 10%of the maximum rate). Orange circles indicates cells with place fields shown in e. **c** Histogram of the Fisher lower bound for decoding error estimated for different positions in the arena. **d** Average firing rate as a function of spatial information. **e** Normalised ratemaps of the 4 cells selected (orange in panels b and d). **f-h** Similar to panel b for for the product (f), DDC (g) and the sampling (h) encoding schemes. **i-m** Same as a-e for experimental session rat 1 day 2. Note that panel j includes putative inhibitory cells (grey) whereas other panels show only putative excitatory neurons.

**Figure S4:**
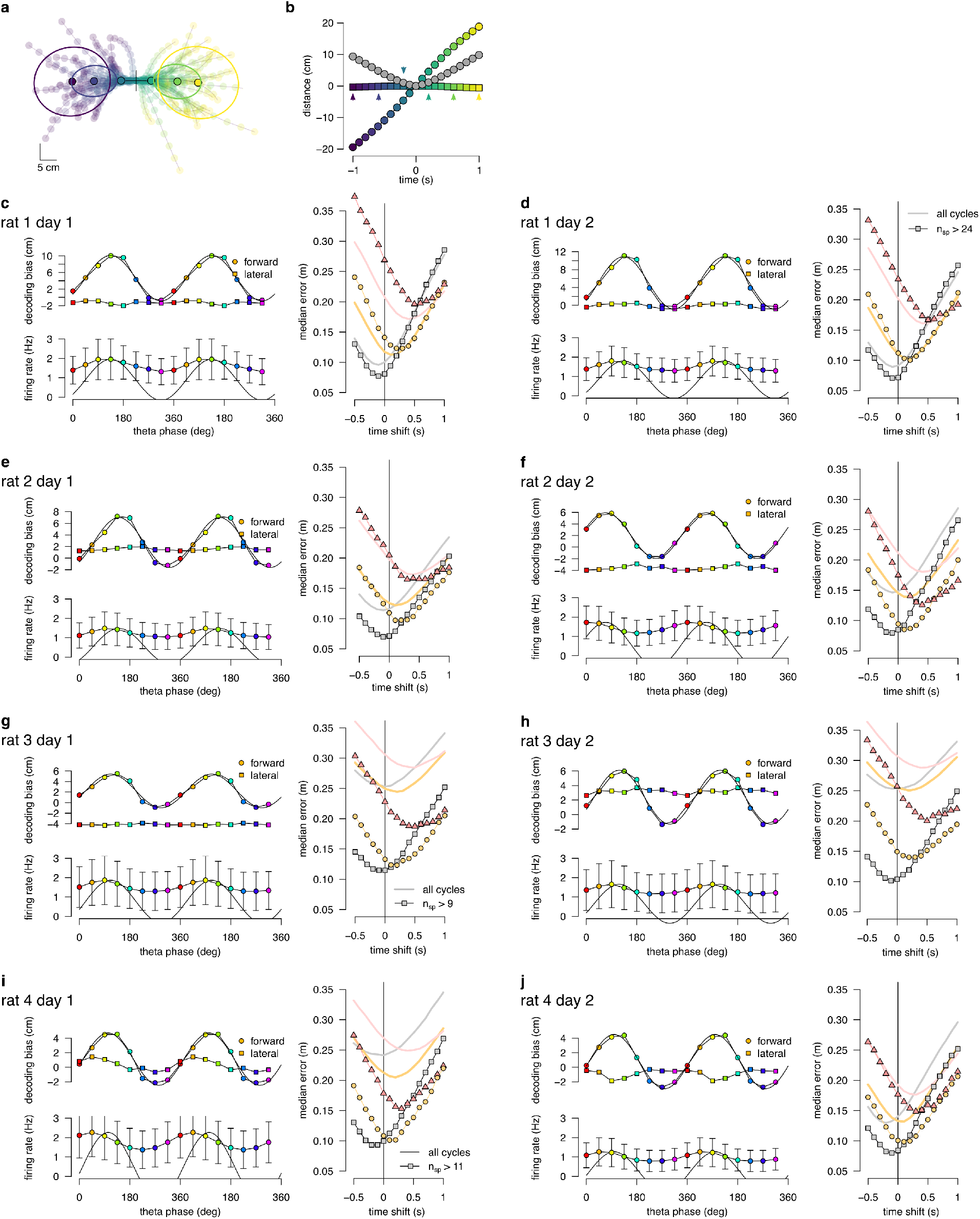
Theta sequence bias and variability in all recording sessions. **a** Location and direction-aligned motion trajectories and 0.5 confidence interval ellipses for −1 ≥ Δ *t* ≥ 1. **b** Bias and spread of motion trajectories as a function of time in an example session. **c** Left: Decoding bias (top) and firing rate as a function of theta phase calculated for all theta cycles. Right: Decoding error for early, mid and late phase spikes calculated as a function of time shift of reference position along the motion trajectory of the animal for all theta cycles (thick lines) or theta cycles with the highest 5% of spike counts (symbols, with the minimum spike count per bin indicated). **d-j** Same as panel c for other recording sessions.

**Figure S5:**
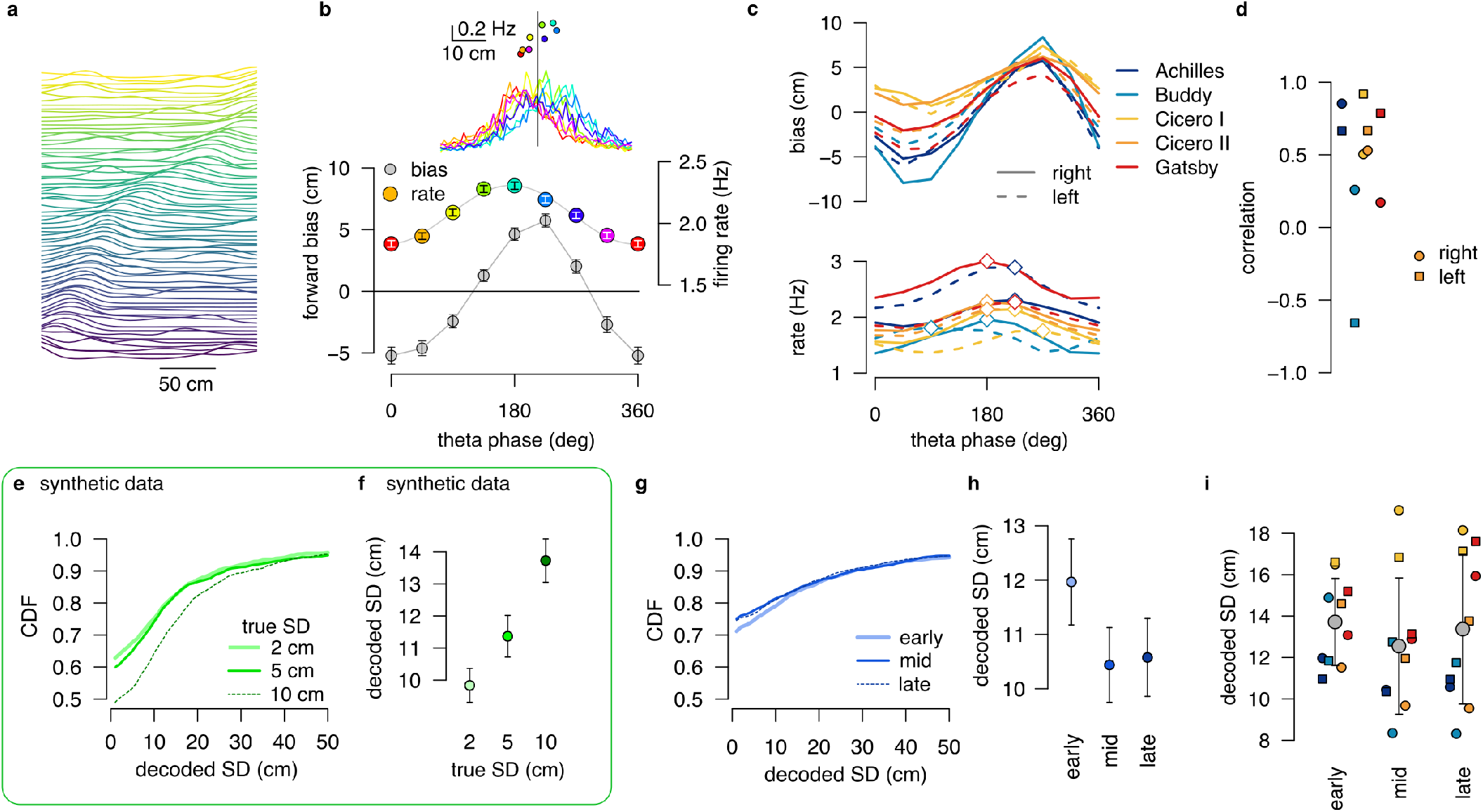
Linear track data and analysis. **a** Normalised firing rate of all putative excitatory neurons recorded in a single session (Achilles, up) order by the location of the peak activity on rightward runs from a previously published dataset (***Grosmark et al., 2016***). **b** Median forward bias and average firing rate as a function of the theta phase calculated in 120^°^windows with 45^°^shift. Inset: the distribution of the decoded location relative to the position of the animal (vertical line) as a function of theta phase (color) and the firing rate as a function of the median decoded location (forward bias, bottom). Error bars show SEM. **c** Median forward bias (top) and average firing rate (bottom) as a function of the theta phase for all recording sessions. Peak forward bias is shifted to 270^°^, diamonds indicate the peak of the firing rate. Color indicate animal name and line style indicates running direction. **d** Correlation between median forward bias and firing rate is typically positive except for one outlier. We note, that the number of recorded neurons was the lowest for the rat Buddy (N=15 cells with place field on right and N=16 on left runs.) **e** Cumulative distribution of the estimated SD for different distributions encoded with a 1 dimensional DDC scheme. Gaussian distributions of different SD were encoded using the tuning functions derived from the place fields shown in A with the same number of theta cycles (N=1846) and bin width (A*r* = 36 ms) as in the experimental data (g-h). **f** Mean decoded SD at different values of the true SD. At this short time window the decoder is still biased (see Fig. 5), but the decoded SD correlates with the true SD. Error bars show SEM. **g** Cumulative distribution of the decoded SD for early, mid and late theta phases for the same session as A-B. We used sharpened tuning functions to reduce the decoding bias. **h** Mean decoded SD at different theta phases. Error bars show SEM. **i** Mean decoded SD at different theta phases for all recorded sessions. Color code is the same as in panel c-d. Only one of the animals (Gatsby) showed an increase in the decoded SD within the theta cycle. Grey symbols show mean and SD across animals. All experimental data presented in this figure was obtained from Grosmark et al. 2016 (ref.14).

**Figure S6:**
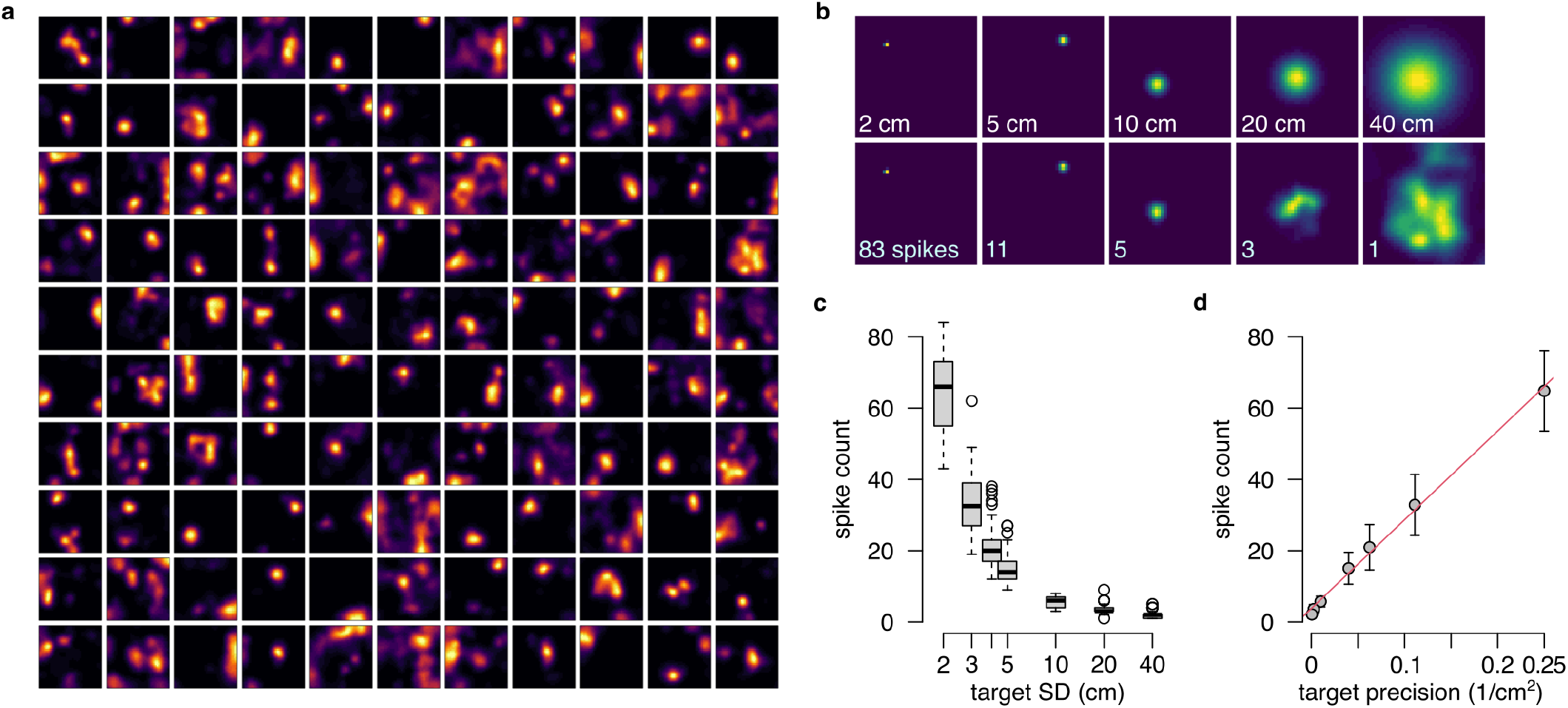
Population gain as a hallmark of the product representation. **a** The 110 tuning curves i//(*x*) from the example session R1D2 used in this analysis as basis function *h(x) =* log *Ψ′(x)* (Eq. 33), with *Ψ′(x) = Ψ(x)/ ∫ Ψ(x) dx*. **b** Representing distributions of increasing uncertainty in the product form using the basis functions shown in panel a. Top: target isotropic Gaussian distributions with increasing SD. Bottom: Approximate distributions with the total number of spikes indicated on the lower left corner. **c** The number of spikes required for optimally approximating a distribution decreases as a function of the target SD. Boxplots show median, quartiles and data range. **d** The number of spikes (mean and SD) as a function of the inverse of the variance (precision) and a linear fit (red) to the data.

**Figure S7:**
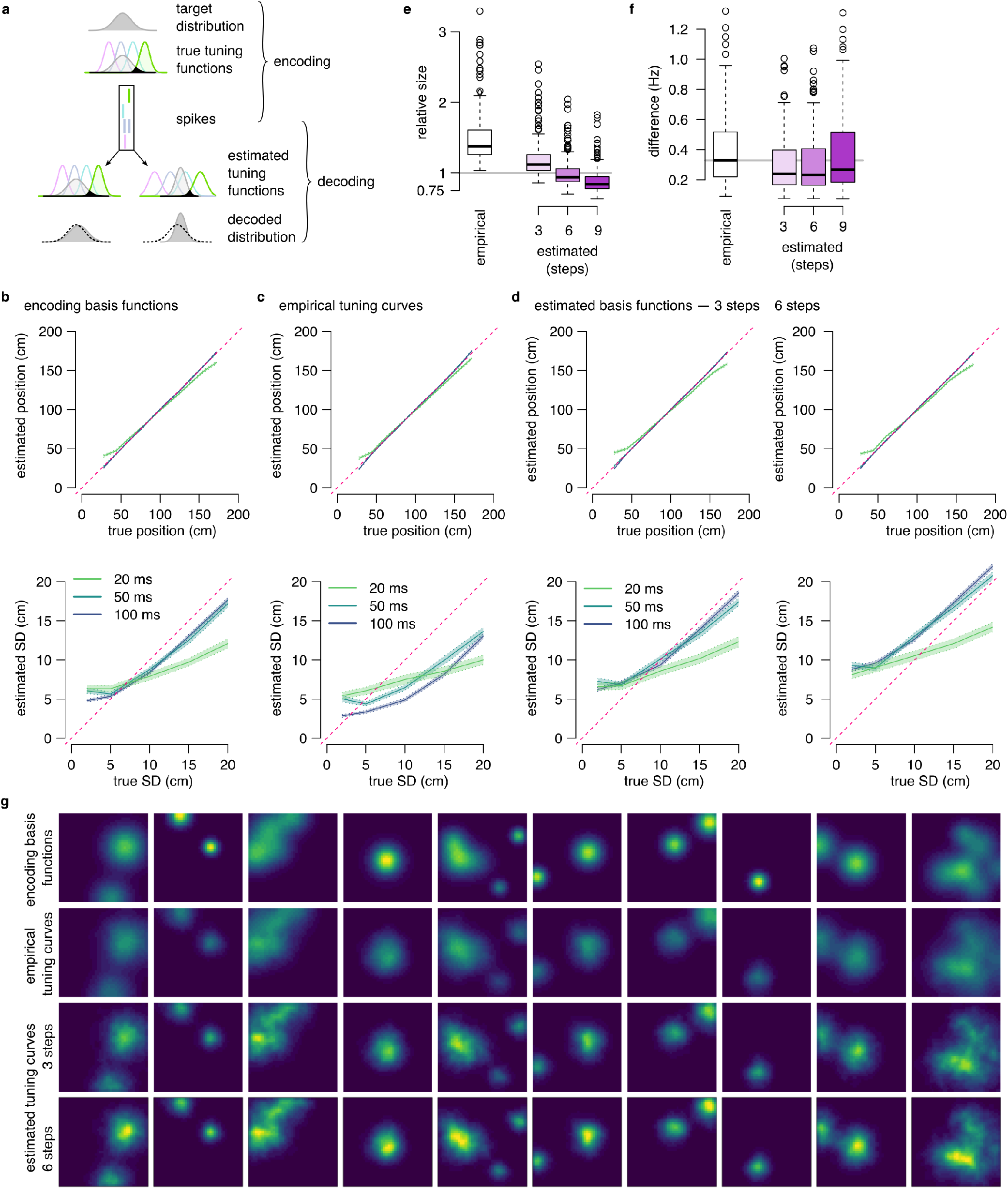
Reducing the bias of the decoded SD in the DDC scheme. **a** Illustration of the problem of decoding bias. In the DDC scheme the firing rate of the neurons is proportional to the overlap between the target distribution and the basis functions of the neurons (black shading). It is possible to approximately reconstruct the encoded distribution from the spikes observed in the population activity and the basis functions (bottom left). However, when the tuning functions used for decoding are different than the basis functions used for encoding than the reconstruction will be biased: Wider tuning functions showing similar overlap with a narrower distribution lead to systematic underestimation of the variance of the encoded distribution (bottom right). **b** Average of the estimated mean (top) and SD (bottom) of 2-dimensional Gaussian distributions decoded in a spike-based DDC scheme with different temporal windows using the true encoding basis functions *ϕ*(*x*). Shading represents the standard error across 1000 repetitions for each combination of time bin and encoded SD. Decoding is approximately unbiased for Δ*t*/*ge*50 ms. **c** Same as b using the empirical rate maps for decoding instead of the true encoding basis functions. The decoded SD is biased even for Δ*t* ≥ /*ge* 50 ms. **d** Same as b using the estimated encoding basis functions after 3 (left) or 6 (right) sharpening steps for decoding. Sharpening the tuning curves for 3 steps eliminates the negative decoding bias while further sharpening introduces a positive bias. **e** Ratio of the estimated and the true encoding basis function sizes for different estimation strategies: empirical tuning curves (place fields) and estimated after 3-9 sharpening steps. The size of the estimated basis function is closest to the original around 6 sharpening steps. **f** Average absolute difference between the true basis function and the estimation across 200 cells is decreased after algorithmic sharpening the tuning curves. The difference is minimal after 3-6 sharpening steps. **g** Examples for the original 2-dimensional basis functions in the synthetic data (top), empirical tuning curves (2nd row) and basis functions estimated using 3 (3rd row) or 6 (bottom) sharpening steps.

**Figure S8:**
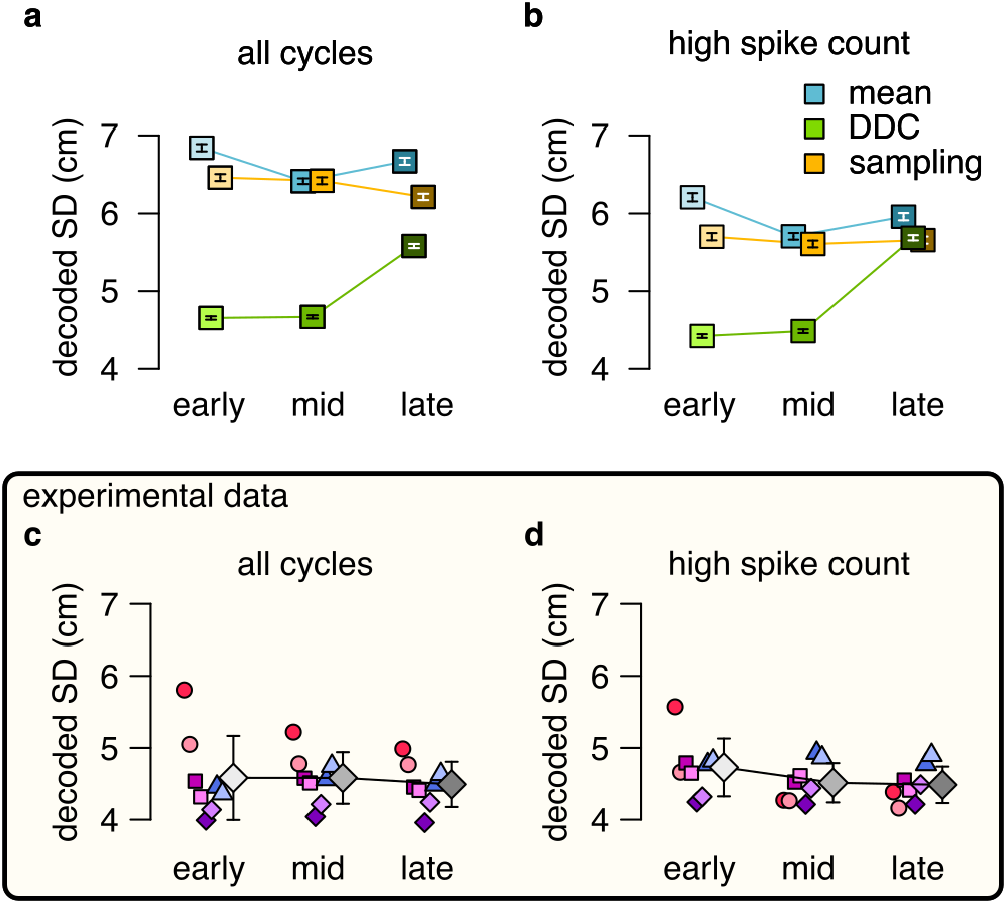
No evidence for DDC code when decoding spikes using the estimated basis functions instead of the empirical tuning curves. **a-b** Estimated standard deviation of the DDC-decoded distribution from spikes at early, mid and late theta phases using the estimated basis functions after 3 sharpening steps using all theta cycles (a) or theta cycles with higher than median spike count (b). Only the distributions encoded via the DDC scheme provides a small, but consistent increase in the estimated SD from early to late theta phase. Symbols show mean and SE. P-values of one-sided, two sample Kolmogorov-Smirnov test comparing the distribution of early versus late for panel a: mean: *P* = 0.99, DDC: *P* = 5 × 10^− 33^, sampling: *P* = 1; for panel b: mean: *P* = 0.99, DDC: *P* = 8.7 × 10^− 28^, sampling: *P* = 0.94. **c-d** Mean of the decoded SD for each animal from early, mid and late theta spikes using all theta cycles (c) or theta cycles with higher than median spike count (d) calculated using the estimated basis functions. Grey symbols show mean and SD across animals. See also Fig. 5 for similar analysis using the empirical tuning curves. P-values of one-sided, two sample Kolmogorov-Smirnov test comparing the distribution of early versus late for panel c: *P >* 0.1 for all animals; panel d: *P >* 0.4 for all animals.

**Figure S9:**
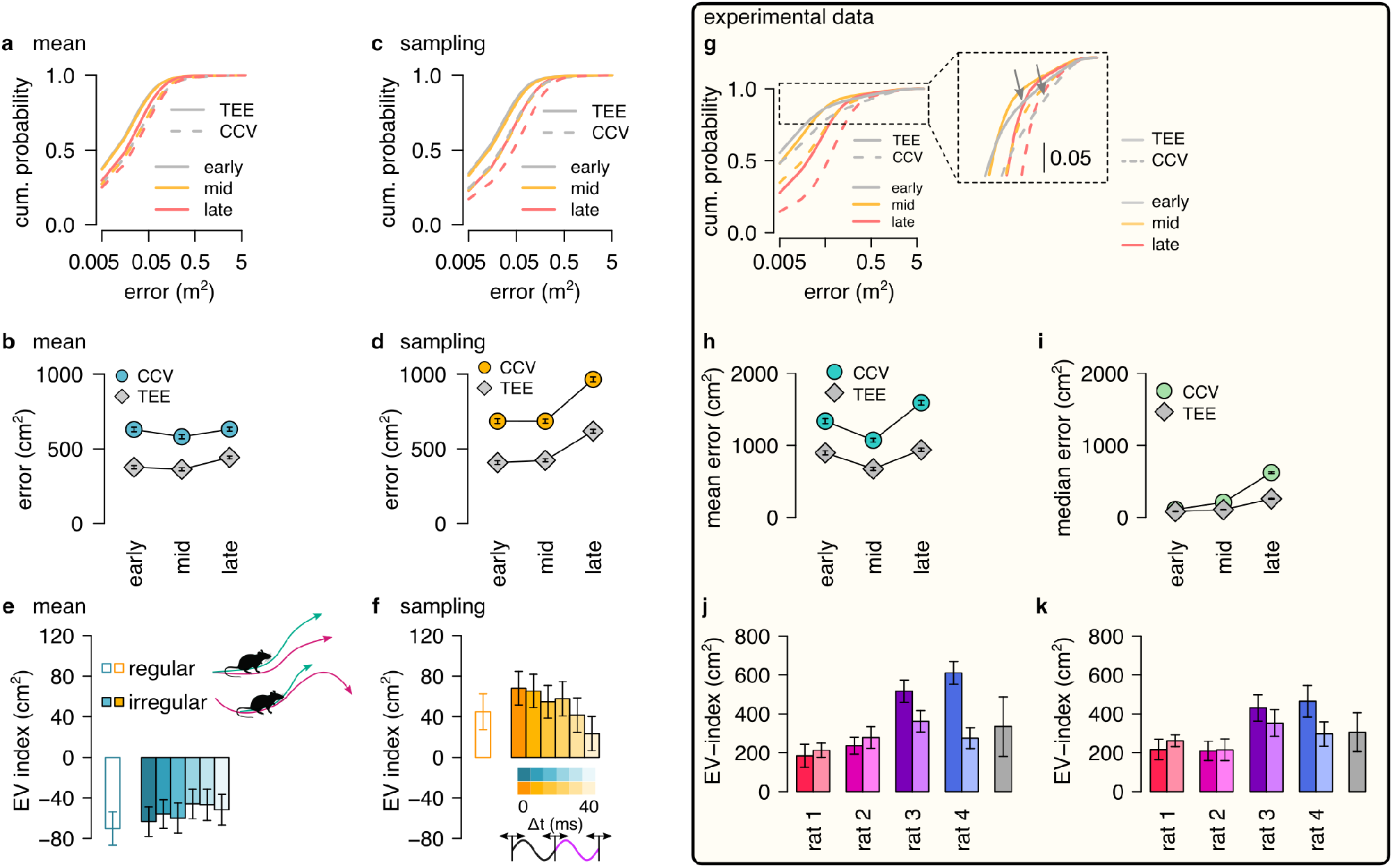
EV-index calculated from all theta cycles. **a** Cumulative distribution of *CCV* (dashed) and *TEE* (solid) for early, mid and late theta phase (colors) in the mean scheme using simulated data. Note the logarithmic x axis. **b** Mean *CCV* and *TEE* calculated from early, mid and late phase spikes in the mean scheme. **c-d** Same as a-b for the sampling scheme. **e-f** The EV-index for the mean (e) and sampling (f) schemes calculated from the mean across theta cycles with simulating regular or irregular theta-trajectories (left inset) and applying various amount of jitter for segmenting the theta cycles (color code, right inset). **g** Cumulative distribution of *CCV* (dashed) and *TEE* (solid) for early, mid and late theta phase (colors) for session R1D2. Note the logarithmic x axis. Arrows in the inset highlight atypically large errors occurring mostly at early theta phase. **h-i** Mean (h) and median (i) of *CCV* and *TEE* calculated from early, mid and late phase spikes for session R1D2. **j-k** EV-index calculated for all analysed sessions (color) and across session mean and SD (grey) using the mean (k) or the median (l) across all theta cycles. See Fig. 6 for the same analysis including only theta cycles with high spike count. Error bars show standard error except for k where they indicate the 5% and 95% confidence interval around the median. P-values for panels e-f and j-k are shown in Table 3.

## References

Bieri KW, Bobbitt KN, Colgin LL. Slow and fast gamma rhythms coordinate different spatial coding modes in hippocampal place cells. Neuron. 2014 May; 82(3):670–81. doi: 10.1016/j.neuron.2014.03.013.

Cei A, Girardeau G, Drieu C, Kanbi KE, Zugaro M. Reversed theta sequences of hippocampal cell assemblies during backward travel. Nature Neuroscience. 2014 May; 17(5):719–724. http://www.nature.com/neuro/journal/v17/n5/full/nn.3698.html, doi: 10.1038/nn.3698.

Colgin LL. Rhythms of the hippocampal network. Nat Rev Neurosci. 2016 Apr; 17(4):239–49. doi: 10.1038/nrn.2016.21.

Colgin LL, Denninger T, Fyhn M, Hafting T, Bonnevie T, Jensen O, Moser MB, Moser EI. Frequency of gamma oscillations routes flow of information in the hippocampus. Nature. 2009 Nov; 462(7271):353–7. doi: 10.1038/nature08573.

Davidson TJ, Kloosterman F, Wilson MA. Hippocampal replay of extended experience. Neuron. 2009 Aug; 63(4):497–507. doi: 10.1016/j.neuron.2009.07.027.

Dayan P, Abbott LF. Theoretical neuroscience. The MIT press; 2001.

Echeveste R, Aitchison L, Hennequin G, Lengyel M. Cortical-like dynamics in recurrent circuits optimized for sampling-based probabilistic inference. Nat Neurosci. 2020 09; 23(9):1138–1149. doi: 10.1038/s41593-020-0671-1.

Fenton AA, Muller RU. Place cell discharge is extremely variable during individual passes of the rat through the firing field. Proc Natl Acad Sci U S A. 1998 Mar; 95(6):3182–7.

Fernández-Ruiz A, Oliva A, Nagy GA, Maurer AP, Berényi A, Buzsáki G. Entorhinal-CA3 Dual-Input Control of Spike Timing in the Hippocampus by Theta-Gamma Coupling. Neuron. 2017 Mar; 93(5):1213–1226.e5. doi: 10.1016/j.neuron.2017.02.017.

Fiser J, Berkes P, Orban G, Lengyel M. Statistically optimal perception and learning: from behavior to neural representations. Trends Cogn Sci. 2010; 14(3):119–30. http://eutils.ncbi.nlm.nih.gov/entrez/eutils/elink.fcgi?cmd=prlinks&dbfrom=pubmed&retmode=ref&id=20153683.

Foster DJ, Wilson MA. Hippocampal theta sequences. Hippocampus. 2007; 17(11):1093–1099. doi: 10.1002/hipo.20345, PMID: 17663452.

Frank LM, Brown EN, Wilson M. Trajectory Encoding in the Hippocampus and Entorhinal Cortex. Neuron. 2000; 27:169–178.

Glimcher P. Decisions, Uncertainty, and the Brain. MIT Press; 2003.

Grosmark AD, Long J, Buzsáki G. Recordings from hippocampal area CA1, PRE, during and POST novel spatial learning. CRCNSorg. 2016; http://dx.doi.org/10.6080/K0862DC5.

Gupta AS, van der Meer MAA, Touretzky DS, Redish AD. Segmentation of spatial experience by hippocampal theta-sequences. Nature neuroscience. 2012 Jul; 15(7):1032–1039. doi: 10.1038/nn.3138.

Hunt LT, Daw ND, Kaanders P, MacIver MA, Mugan U, Procyk E, Redish AD, Russo E, Scholl J, Stachenfeld K, Wilson CRE, Kolling N. Formalizing planning and information search in naturalistic decision-making. Nat Neurosci. 2021 Jun; doi: 10.1038/s41593-021-00866-w.

Jezek K, Henriksen E, Treves A, Moser E, Moser MB. Theta-paced flickering between place-cell maps in the hippocampus. Nature. 2011 Oct; 478(7368):246–249. doi: 10.1038/nature10439.

Johnson A, Redish AD. Neural ensembles in CA3 transiently encode paths forward of the animal at a decision point. J Neurosci. 2007 Nov; 27(45):12176–89. doi: 10.1523/JNEUROSCI.3761-07.2007.

Kao TC, Sadabadi MS, Hennequin G. Optimal anticipatory control as a theory of motor preparation: A thalamocortical circuit model. Neuron. 2021 May; 109(9):1567–1581.e12. doi: 10.1016/j.neuron.2021.03.009.

Kay K, Chung JE, Sosa M, Schor JS, Karlsson MP, Larkin MC, Liu DF, Frank LM. Constant Sub-second Cycling between Representations of Possible Futures in the Hippocampus. Cell. 2020 02; 180(3):552–567.e25. doi: 10.1016/j.cell.2020.01.014.

Kelemen E, Fenton AA. Dynamic grouping of hippocampal neural activity during cognitive control of two spatial frames. PLoS Biol. 2010 Jun; 8(6):e1000403. doi: 10.1371/journal.pbio.1000403.

Kutschireiter A, Surace SC, Sprekeler H, Pfister JP. Nonlinear Bayesian filtering and learning: a neuronal dynamics for perception. Sci Rep. 2017 08; 7(1):8722. doi: 10.1038/s41598-017-06519-y.

Lange RD, Shivkumar S, Chattoraj A, Haefner RM. Bayesian Encoding and Decoding as Distinct Perspectives on Neural Coding. bioRxiv. 2020; (https://doi.org/10.1101/2020.10.14.339770).

Ma WJ M BJ,, Latham PE, Pouget A. Bayesian inference with probabilistic population codes. Nat Neurosci. 2006; 9:1432–1438.

MacKay DJC. Information Theory, Inference, and Learning Algorithms. Cambridge University Press; 2003.

Mante V, Sussillo D, Shenoy KV, Newsome WT. Context-dependent computation by recurrent dynamics in prefrontal cortex. Nature. 2013 Nov; 503(7474):78–84. http://www.nature.com/nature/journal/v503/n7474/abs/nature12742.html, doi: 10.1038/nature12742.

Mastrogiuseppe F, Ostojic S. Linking Connectivity, Dynamics, and Computations in Low-Rank Recurrent Neural Networks. Neuron. 2018 08; 99(3):609–623.e29. doi: 10.1016/j.neuron.2018.07.003.

Mattar MG, Daw ND. Prioritized memory access explains planning and hippocampal replay. Nat Neurosci. 2018 11; 21(11):1609–1617. doi: 10.1038/s41593-018-0232-z.

Miller KJ, Botvinick MM, Brody CD. Dorsal hippocampus contributes to model-based planning. Nat Neurosci. 2017 Sep; 20(9):1269–1276. doi: 10.1038/nn.4613.

Murphy KP. Machine Learning: A Probabilistic Perspective. MIT Press, Cambridge, MA; 2012.

Murray I. Advances in Markov chain Monte Carlo methods. PhD thesis, UCL; 2007.

Neftci EO, Averbeck BB. Reinforcement learning in artificial and biological systems. Nature Machine Intelligence. 2019; 1:133–143.

O’Keefe J, Nadel L. The Hippocampus as a Cognitive Map. Oxford University Press; 1978. http://www.cognitivemap.net/.

Orbán G, Berkes P, Fiser J, Lengyel M. Neural Variability and Sampling-Based Probabilistic Representations in the Visual Cortex. Neuron. 2016 Oct; 92(2):530–543. doi: 10.1016/j.neuron.2016.09.038.

Pfeiffer BE, Foster DJ. Hippocampal place-cell sequences depict future paths to remembered goals. Nature. 2013 May; 497(7447):74–79. http://www.nature.com/nature/journal/v497/n7447/full/nature12112.html?WT.ec_id=NATURE-20130502, doi: 10.1038/nature12112.

Pouget A, Beck JM, Ma WJ, Latham PE. Probabilistic brains: knowns and unknowns. Nat Neurosci. 2013 Sep; 16(9):1170–8. doi: 10.1038/nn.3495.

Redish AD. Vicarious trial and error. Nat Rev Neurosci. 2016 Mar; 17(3):147–59. doi: 10.1038/nrn.2015.30.

Rich PD, Liaw HP, Lee AK. Large environments reveal the statistical structure governing hippocampal representations. Science. 2014 Aug; 345(6198):814–7. doi: 10.1126/science.1255635.

Robbe D, Montgomery SM, Thome A, Rueda-Orozco PE, McNaughton BL, Buzsaki G. Cannabinoids reveal importance of spike timing coordination in hippocampal function. Nat Neurosci. 2006 Dec; 9(12):1526–33. doi: 10.1038/nn1801.

Savin C, Deneve S. Spatio-temporal Representations of Uncertainty in Spiking Neural Networks. In: Proceedings of the 27th International Conference on Neural Information Processing Systems; 2014..

Schomburg EW, Fernández-Ruiz A, Mizuseki K, Berényi A, Anastassiou CA, Koch C, Buzsáki G. Theta phase segregation of input-specific gamma patterns in entorhinal-hippocampal networks. Neuron. 2014 Oct; 84(2):470–85. doi: 10.1016/j.neuron.2014.08.051.

Sezener CE, Dezfouli A, Keramati M. Optimizing the depth and the direction of prospective planning using information values. PLoS Comput Biol. 2019 03; 15(3):e1006827. doi: 10.1371/journal.pcbi.1006827.

Skaggs WE, McNaughton BL, Wilson MA, Barnes CA. Theta phase precession in hippocampal neuronal populations and the compression of temporal sequences. Hippocampus. 1996; 6(2):149–72. http://eutils.ncbi.nlm.nih.gov/entrez/eutils/elink.fcgi?cmd=prlinks&dbfrom=pubmed&retmode=ref&id=8797016.

Stachenfeld KL, Botvinick MM, Gershman SJ. The hippocampus as a predictive map. Nat Neurosci. 2017 Nov; 20(11):1643–1653. doi: 10.1038/nn.4650.

Stroud JP, Porter MA, Hennequin G, Vogels TP. Motor primitives in space and time via targeted gain modulation in cortical networks. Nat Neurosci. 2018 12; 21(12):1774–1783. doi: 10.1038/s41593-018-0276-0.

Tang W, Shin JD, Jadhav SP. Multiple time-scales of decision-making in the hippocampus and prefrontal cortex. Elife. 2021 Mar; 10. doi: 10.7554/eLife.66227.

Ujfalussy BB, Makara JK, Branco T, Lengyel M. Dendritic nonlinearities are tuned for efficient spike-based computations in cortical circuits. Elife. 2015 Dec; 4. doi: 10.7554/eLife.10056.

Vértes E, Sahani M. Flexible and accurate inference and learning for deep generative models. Advances in Neural Information Processing Systems. 2018; p. 4166–4175.

Vértes E, Sahani M. A neurally plausible model learns successor representations in partially observable environments. In: NeurIPS; 2019..

Vyas S, Golub MD, Sussillo D, Shenoy KV. Computation Through Neural Population Dynamics. Annu Rev Neurosci. 2020 07; 43:249–275. doi: 10.1146/annurev-neuro-092619-094115.

Wainwright MJ, Jordan MI. Graphical Models, Exponential Families, and Variational Inference. Foundations and Trends in Machine Learning. 2008; 1(1-2):1–305.

Walker EY, Cotton RJ, Ma WJ, Tolias AS. A neural basis of probabilistic computation in visual cortex. Nat Neurosci. 2020 01; 23(1):122–129. doi: 10.1038/s41593-019-0554-5.

Wang M, Foster DJ, Pfeiffer BE. Alternating sequences of future and past behavior encoded within hippocampal theta oscillations. Science. 2020 10; 370(6513):247–250. doi: 10.1126/science.abb4151.

Wikenheiser AM, Redish AD. Hippocampal theta sequences reflect current goals. Nat Neurosci. 2015 Feb; 18(2):289–94. doi: 10.1038/nn.3909.

Wilson MA, McNaughton BL. Dynamics of the hippocampal ensemble code for space. Science. 1993; 261:1055–1058.

Zemel R, Dayan P, Pouget A. Probabilistic interpretation of population codes. Neural Comput. 1998; 10:403–430.

Zhang K, Ginzburg I, McNaughton BL, Sejnowski TJ. Interpreting neuronal population activity by reconstruction: unified framework with application to hippocampal place cells. J Neurophysiol. 1998 Feb; 79(2):1017–44. doi: 10.1152/jn.1998.79.2.1017.

Zheng C, Hwaun E, Lee Colgin L. Impairments in Hippocampal Place Cell Sequences during Errors in Spatial Memory. bioRxiv. 2020; https://doi.org/10.1101/2020.04.20.051755.

